# The Endothelial Cell-Expressed Prion Protein Prnd/Doppel Promotes Neovascularization and Long-term Recovery after Ischemic Stroke

**DOI:** 10.64898/2026.02.21.707233

**Authors:** Tao Wang, Sehee Kim, John E. Morales, Joseph H. McCarty

**Author notes:** Corresponding Author: Joseph H. McCarty Department of Neurosurgery, Unit 1004 University of Texas M. D. Anderson Cancer enter 1515 Holcombe Boulevard Houston, TX 77030.

## Abstract

Ischemic stroke remains a leading cause of mortality and long-term disability, yet therapeutic options for promoting recovery remain severely limited. Here, we investigate the role of Prnd/Doppel, a prion family member, in stroke pathophysiology and recovery. Using middle cerebral artery occlusion (MCAO) in mice genetically null for Prnd (KO), inducibly overexpressing Prnd in endothelial cells (ECs), or wild-type (WT) controls, we assessed outcomes through infarct volume measurements, behavioral analysis, and immunohistochemical evaluation of vascular integrity and inflammation. While acute infarct volumes at 24 hours were comparable between WT and KO mice, striking differences emerged during recovery: KO mice exhibited significantly impaired functional outcomes at both 14 and 30 days post-MCAO, accompanied by disorganized cerebrovascular architecture, increased brain atrophy, and elevated CD68-positive inflammatory infiltration by day 30. Conversely, endothelial-specific Prnd overexpression, though not affecting acute outcomes, markedly enhanced tight junction protein expression at day 7, promoted angiogenesis, and improved long-term neuronal survival in the ischemic territory. These findings establish Prnd as a critical mediator of post-stroke vascular remodeling and functional recovery, distinguishing it from acute neuroprotective mechanisms. Our results identify Prnd as a promising therapeutic target for enhancing organized neovascularization and promoting sustained functional recovery following ischemic stroke, with potential applications to other neurological disorders characterized by cerebrovascular dysfunction.

## Introduction

As the body’s most densely vascularized organ, the brain relies on a highly specialized network of blood vessels to deliver the oxygen and nutrients required to sustain its metabolic activity [1]. Ischemic stroke impairs cerebral blood flow, disrupts oxygen delivery and energy metabolism, which leads to cognitive and motor deficits and increases the risk of long-term complications including dementia [2, 3]. The neurovascular unit (NVU) is a multicellular structure composed of neurons, glia (astrocytes and microglia), vascular mural cells (vSMCs), pericytes (PCs), ECs, and oligodendrocytes [4]. These cellular components work together to maintain brain homeostasis through regulation of blood-brain barrier (BBB) permeability and cerebral blood flow (CBF) [5]. During the acute phase after stroke onset, NVU dysfunction contributes to decreased tight junction integrity which increases BBB permeability, elevating the risk of vasogenic cerebral edema [6, 7]. Consequently, NVU repair is critical for preventing adverse outcomes following stroke and may offer novel therapeutic strategies for stroke management [8, 9].

The Prnd gene encodes the Doppel protein, which is a member of the prion protein family which also includes the canonical cellular prion protein PrPc [10], which is highly expressed in the neurons of the adult brain [11]. In contrast, Prnd is specifically expressed in angiogenic ECs during development [12]. PrPc expression persists in adulthood, whereas Prnd expression is greatly reduced in the quiescent adult brain vasculature [12]. PrPc plays a role in protecting neurons after ischemic stroke, with PrPc deficiency leading to worsened brain damage and outcome [13]. Prnd plays an important role in developmental angiogenesis and formation of the BBB [12, 14]. Prnd mRNA is expressed in vascular ECs of angiogenic blood vessels in the developing brain, with expression levels peaking early postnatally and decreasing in the adult brain once vascular proliferation and sprouting cease and blood vessel display a quiescent status [14]. In our previous studies, we reported that Prnd deficiency induces abnormal embryonic angiogenesis and impaired BBB formation in the neonatal brain [12]. Biochemical assays and primary EC culture systems reveal that GPI-linked Doppel activates transmembrane receptor tyrosine kinases with central roles in blood vessel survival, metabolism and migration [12]. While these previous results revealed important functions for Prnd in CNS blood vessel morphogenesis and maturation and suggested that targeting Prnd to promote or block angiogenesis could be an effective therapeutic approach to treat vascular-related CNS pathologies, the role of the Prnd in ischemic stroke remains largely unknown.

In this study, we analyzed the functions of Prnd in ischemic stroke using complementary genetic mouse models, including Prnd loss of function knockout mice (Prnd-/-) as well as a novel gain of function model in which Prnd is inducibly expressed in vascular ECs. Our results demonstrate that Prnd significantly promotes angiogenesis, facilitates BBB repair, and enhances long-term neuronal survival following ischemic injury. Furthermore, we provide evidence that modulation of Prnd expression directly influences NVU integrity and functional recovery after stroke. Collectively, our data strongly suggest that Prnd represents a novel and promising therapeutic target with translational potential to promote long-term recovery not only in brain ischemia but possibly in other neurological diseases characterized by neurovascular dysfunction and impaired tissue repair mechanisms.

## Materials and Methods

### Experimental mice

All mice were housed in the animal care facility at the MD Anderson Cancer Center at Texas University, which is approved by the Institutional Animal Care and Use Committee (IACUC) and the MD Anderson Subcommittee on Animal Studies, both of which are AAALAC-accredited. Mice were housed under controlled environmental conditions (12h light:12h darkness) and acclimated for at least 1 week. For MCAO surgery, 10–14 weeks old aged matched male mice were used. All animal experiments were conducted in a blind fashion. Generation of the Prnd-/- mouse model has been described [15]. To generate the Cre-inducible Prnd overexpression model, a targeting cassette containing a CAG promoter (pCAG) and loxSTOPlox sequence upstream of the mouse Prnd cDNA was inserted in the Rosa26 locus of mouse iTL BA1 (129/SvEv x C57BL/6) hybrid embryonic stem cells. Cells were microinjected into C57BL/6 blastocysts. Resulting chimeras with a high percentage agouti coat color were mated to wild-type C57BL/6N mice to generate F1 heterozygous offspring. Tail DNA was analyzed as described below from pups with agouti or black coat color. Standard loxP-STOP-loxP cassette design for ROSA26 knock in contains the Neomycin selection marker within that floxed STOP cassette, and the Neo-STOP component is removed during the Cre-loxP recombination.

### Brain ischemia

Transient focal ischemia was induced by suture occlusion of the middle cerebral artery in male mice anaesthetized using 1.5% isofluorane, 70% N2O and 30% O2. Ischemia was induced by introducing a coated filament (702223; Doccol) from the external carotid artery into the internal carotid artery and advancing it into the circle of Willis to the branching point of the left MCA, thereby occluding the middle cerebral artery [16, 17]. Rectal temperature was maintained at 37°C with a thermostatically controlled heating pad. All surgical procedures were performed under an operating stereomicroscope. The causes of animal exclusion as we have used earlier [16] include: 1) Mice that showed no neurological deficits. 0-2 mice were excluded in each group due to this reason. 2) Brains with evidence of surgical subarachnoid hemorrhage. 0-1 mice were excluded in each group due to surgical subarachnoid hemorrhage. Mice with secondary hemorrhage after stroke were not excluded.

24 hours after MCAO, mice were sacrificed and brains were sectioned coronally at 1 mm intervals and stained by vital dye immersion: (2%) 2,3,5-triphenyltetrazolium hydrochloride (TTC) to quantify infarct volume. 24 hours after MCAO, the mice were deeply anesthetized. The brains were rapidly removed and coronally sectioned into 7 slices with 1 mm thickness, stained in 2% TTC (Sigma, St. Louis, MO) at 37 °C for 15 min, and fixed in 4% paraformaldehyde for 24 h [18]. The brain slices were scanned, and infarctions were measured using NIH ImageJ software as we have used earlier [17]. The infarction in each section was normalized to the non-ischemic contralateral side and expressed as a percentage of the contralateral hemisphere with the following formula: Infarct size = (contralateral area - ipsilateral non-infarct area) / contralateral area × 100%.

Brain atrophy at 13 or 28 days after stroke was measured by Nissl staining [16]. Mice were euthanized at either 13 or 28 days after ischemia and then perfused with ice-cold PBS followed with 4% (w/v) paraformaldehyde in PBS. Using a vibratome (Leica VT1000, Wetzlar, Germany), 50 μm coronal brain slices were cut after post-fixation with 4% (w/v) paraformaldehyde for 24 h. Brain slices were stained in 0.1% cresyl violet solution for 30 min and then rinsed in distilled water. Stained sections were fixed by serial dehydration in ethanol (70%, 90%, 100%) and xylene. Fixed slices were then scanned and quantified with NIH ImageJ software, as mentioned above [16]. The brain atrophy in each section was normalized to the non-ischemic contralateral side and expressed as a percentage of the contralateral hemisphere with the following formula: Brain atrophy = (contralateral area - ipsilateral area) / contralateral area × 100%.

### Neurological deficit measurements

For all animals, the neurological deficit scores were evaluated based on the established methods [19]. The scores for the neurological behavioral were graded from 0 to 5 and as follows: 0= no observed deficits, 1= forelimb flexion when lifted by the tail, 2= consistently reduced resistance to lateral push, 3= unilateral circling toward the paretic side, 4= ambulation inability or difficulty, 5= dead (normal, 0; maximal deficit score, 5).

### Isolation of CD31^+^ primary mouse brain ECs

Mouse were euthanized by CO_2_ in accordance with IACUC regulations. Mouse brains were dissected, manually dissociated and CD31-positive cells selected using the Miltenyi Biotec Adult Brain Dissociation Kit (130-107-677), CD31 Microbeads (130-097-418) and LS Columns (130-042-401). Briefly, the brains were minced, enzymatically digested and homogenized by means of gentle trituration. The brain homogenate was filtered through a 70 µm cell strainer to obtain a single cell suspension. Debris and myelin were removed by phase separation using the debris removal solution contained in the kit. Red blood cells were removed by lysis using the red blood cell removal solution. The remaining cells were positive selected for expressing mouse CD31 antigen by labeling with Miltenyi Biotec CD31 Microbeads and then magnetically separated with Miltenyi Biotec LS Columns. The final eluted CD31-positive cell fraction was washed twice in PBS and either flash frozen on dry ice or cultured on laminin-coated plates in DMEM-F12 medium supplemented with 5% FBS, 1% ECGS and penicillin-streptomycin.

### Human brain EC experiments

Primary HBMECs were purchased from ScienCell Research Laboratories (cat#1000), and were maintained in EC medium (ECM, cat#1001). Cells were grown on dishes coated with rat tail collagen (Sigma-Aldrich) at 37°C and 5% CO2. For lentivirus infections, HBMECs were seeded into six-well plates at a concentration of 5×10^5^ cells/well and cultured in complete medium for 24 h. Cells were infected with pLOC lentivirus expressing human PRND or with lentivirus-mediated pLOC negative control (GFP/RFP). After 48 h, cells are collected for Western Blot analysis. The cell lysate was prepared using RIPA lysis buffer.

### RNA extraction and Polymerase Chain Reaction (PCR)

Total RNA was isolated using Tri reagent (MRC, OH, USA). The RNA was then precipitated with isopropanol, and the pellet was washed with 70% ethanol, air-dried, and dissolved in sterile diethylpyrocarbonate (DEPC) water. The concentration and purity of RNA were determined with a NanoDrop spectrophotometer (NanoDrop Technologies, Inc., Wilmington, DE, USA). 1µg of total RNA was reverse transcribed using reverse-transcribed using oligo (dT) primers and Superscript II reverse transcriptase (Life Technologies, Inc.) according to the manufacturer’s protocol. The PCR reaction was performed using the SYBR Green I (Applied Biosystems, Foster City, CA, United States) and specific PCR primers for Cdh5/VE-Cadherin, KDR/VEGFR2, Prnd, GAPDH, and Real-Time PCR system (Applied Biosystems). Standard PCR was performed using 2x Taq supermix (Syd Labs). Cycling condition was 95°C 4’; 35 cycles of (94° C 20”,53° C 20”, 72°C 20”); 72° C 2’;12°C hold.

### Immunoblot analyses

Mice brain tissues or cells were homogenized and lysed in a buffer containing 10M Urea and 1% SDS. Lysates were separated by 12% SDS-PAGE and electro transferred to a polyvinylidene difluoride membrane (Bio-Rad). The membranes were soaked in blocking buffer (1× Tris-buffered saline, 1% Tween 20 and 5% nonfat dry milk) for 1 h and incubated overnight at 4°C with the primary antibodies against Prnd [12]. Blots were washed three times in TBS-T buffer and developed using a peroxidase-conjugated anti-mouse IgG and a western blotting detection kit (Amersham;). The bands were visualized using a ChemicDoc (Bio-Rad) and quantified using Image J software (National Institutes of Health, Bethesda, MD).

### Doppel antibody generation

Anti-Doppel polyclonal rabbit polyclonal antibodies were generated using a synthetic 20-residue peptide sequence corresponding to amino acids 55-73 (RPGAFIKQGRKLDIDFGAEC) of the human Doppel protein. This sequence is 100% identical to the mouse Doppel protein, as determined by sequence alignment using BLAST. Pre-bleeds were taken from two rabbits in parallel with synthesis of the selected peptide conjugated to the immunogenic KLH carrier protein. Six weeks later, the rabbits were immunized by subcutaneous injection with the KLH conjugated peptide. After 10 days, the first production bleed was taken from each rabbit. 10 days after the bleed, the rabbits were delivered a second booster immunization. After another 10 days, the rabbits were bled a second time. Antibody specificities from each bleed were tested by ELISA, immunoblotting and immunohistochemistry. Promising bleeds were affinity purified using the immunizing peptide. This antibody was only used in Western Blot.

### Frozen Sectioning and Immunostaining

The mice were perfused with phosphate buffered saline (PBS) followed by 30 ml of 4% paraformaldehyde in PBS. The brains were then isolated and post fixation in 2% paraformaldehyde for one night. The brains were protected overnight in 30% sucrose, and sectioned coronally at 20 µm thickness. The slices were adhered to SuperFrost plus glass slides, air-dried overnight, and stored at - 20°C until use. For immunostaining, the slices were rehydrated and permeabilized with 1% Triton in PBS for 15min. The sections were blocked sequentially (20min each at room temperature) with 50 mM NH4Cl and blocking buffer (PBS, 10% Horse serum, 5 mg/ml BSA, 0.2% Triton X-100). For antibody dilution and washing, the buffer was a 1:4 dilution of the blocking buffer with PBS. We added primary antibodies: Prnd (Thermo Fisher Scientific, PIPA5113502), VEGF (Thermo Fisher Scientific, ENP802), CD31 (R&D Systems, AF3628), ZO-1 (Thermo Fisher Scientific, PIPA585256), Claudin 5 (Thermo Fisher Scientific, 35-2500), GFAP (Novus, NBP1-05198), CD68 (Novus, NBP2-33337), Neun (Thermo Fisher Scientific, PIPA5143552), TMEM 119 (Novus Biologicals, NBP230551), Laminin (Sigma, L9393), and VE-Cadherin (BD Pharmingen, 550548) to the slides and incubated the slides overnight at 4°C in a moisturized dark box, washed three times (5min per wash with gentle rocking at room temperature) and added secondary antibodies (1:500 dilutions) and incubated at room temperature for 1 hr. The slides were washed another three times. The slides were mounted with either 70% glycerol or DAPI mounting media (Vector Laboratories), kept at 4°C until imaging. Confocal images were acquired using an Olympus Fluoview FV3000 4×, 20× objectives. All comparative images were taken with the same laser power and gain settings to make qualitative comparisons between staining levels in different samples.

Immunofluorescence images were acquired using Olympus Fluoview FV3000 confocal laser scanning microscope. Multidimensional acquisition was conducted using Z stacks with 2 mm slicing intervals at a scan rate of 4 μs/pixel with a resolution of 1024 × 1024 pixels per slice and digitally compiled in FV31S-SW (version 2.4.1.198). Image acquisition parameters, including exposure time, laser power, gain, and voltages, were fixed for the compared imaging channel. All images were analyzed using ImageJ. Serial Z-stack images (3 slices per image) were used for quantitation, and all images were scaled (µm) as per the objective lens used for acquisition. Image stacks were projected for maximum intensity to include all signals.

### Quantitative analysis of blood vessels

Mice brain slices were stained with CD31. The images were captured by Olympus Fluoview FV3000 confocal laser scanning microscope and were analyzed by AngioTool for quantitative analysis of vascular networks [20]. AngioTool is open source and can be downloaded at http://angiotool.nci.nih.gov. Several morphometric parameters are computed including total and average vessel length, branching point density, and vascular density. Some metrics are normalized to the area of the convex hull containing the region covered by the vessels allowing for comparison of differently sized vascular networks. Additionally, a fast box counting algorithm has been implemented for computation of lacunarity, an index for vascular structural nonuniformity, which is reported as the average lacunarity over all box sizes.

### Statistical analysis

Values are expressed as mean with SD. Statistical analysis was performed with GraphPad Prism 10.3.1 software using one-way ANOVA or unpaired t test. Statistical significance was accepted at the 95% confidence level (P < 0.05).

## Results

### Prnd deficiency impairs adult cerebral blood vessel morphogenesis

To investigate the role of Prnd in cerebrovascular development and homeostasis, we used AngioTool to perform quantitative morphometric analysis of brain vascular networks in 12-week-old Prnd WT and Prnd KO mice using anti-CD31 immunofluorescence imaging [15]. We assessed multiple vascular parameters including vessel area, percentage of vessels per area, total vessel length, and average vessel length. Our analysis revealed significant reductions in most of the measured vascular parameters in Prnd KO mice compared to WT controls (Fig. 1A, B). These findings demonstrate that genetic ablation of Prnd leads to impaired cerebrovascular morphogenesis and maturation in adult mice. This observation is consistent with our previous findings that Prnd plays a critical role in developmental angiogenesis and BBB formation [12].

**Figure 1.**
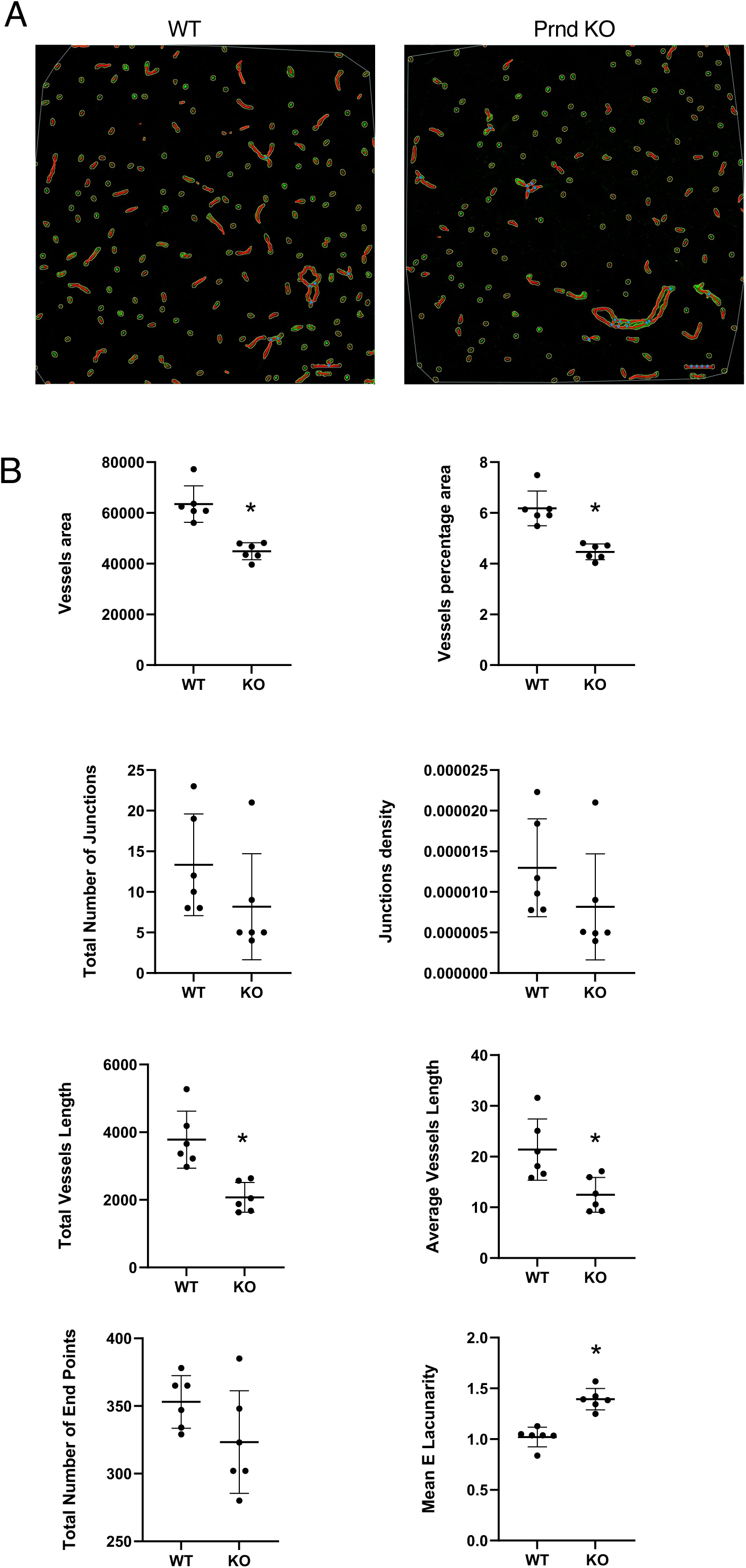
Prnd Deficiency Impairs Cerebrovascular Morphogenesis and Maturation in Adult Mice. (**A**); Representative immunofluorescence images from coronal brain sections through the striatum of 12-week-old Prnd WT and KO mice. Sections were stained with anti-CD31 antibodies to visualize vascular endothelial cells. Images were pseudo-colored using AngioTool software for quantitative vascular network analysis. Scale bars represent 50 μm. **(B);** Quantitative morphometric analysis of vascular networks in the striatum of 12-week-old Prnd WT and KO mice. All cerebral blood vessel parameters are significantly reduced in Prnd KO mice compared to WT controls, indicating impaired cerebrovascular development and maturation (n = 6 mice per group). Data are presented as means ± SEM; *p < 0.05 using two-tailed Student’s t-test.

### Prnd deficiency has no effect on short-term but worsened long-term ischemic stroke outcome

To determine whether Prnd deficiency influences stroke pathology, we evaluated brain infarct volumes in Prnd WT and KO mice at multiple time points (24 hours, 3 days, 13 days, and 28 days) following MCAO. During the acute phase (24 hours and 3 days post-stroke), infarct volumes were comparable between Prnd WT and KO mice. (Fig. 2A, 2B). However, striking differences emerged during the chronic recovery phase. At 13 days and 28 days post-MCAO, Prnd KO mice exhibited significantly greater brain atrophy (loss of brain volume) compared to WT controls (Fig. 2B).

**Figure 2.**
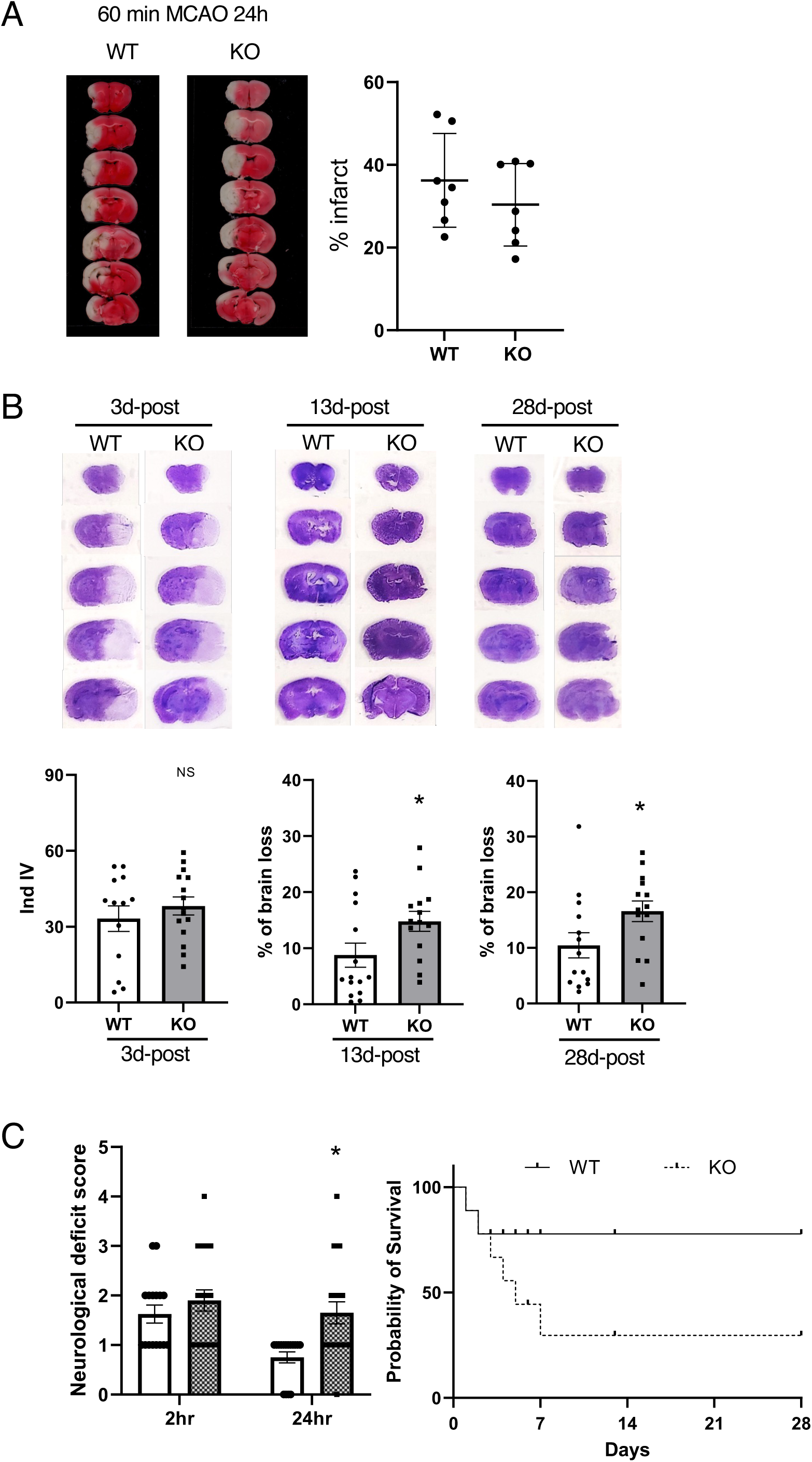
Prnd Deficiency Has No Effect on Acute Stroke Injury but Exacerbates Long-Term Outcomes. **(A);** Assessment of acute brain injury in Prnd WT and KO mice at 24 hours following MCAO. Mice were subjected to 60 minutes of focal cerebral ischemia, and infarct volumes were quantified in TTC-stained coronal brain sections. Infarct volumes are comparable between genotypes at this acute time point (n = 7 Prnd WT, n = 7 Prnd KO). Data are presented as means ± SEM using two-tailed Student’s t-test. **(B);** Time-course analysis of brain injury in Prnd WT and KO mice following ischemic stroke. Mice were subjected to 30 minutes of MCAO, and both infarct volume and total brain volume were measured in Nissl-stained coronal sections at 3 days (n = 13 Prnd WT, n = 15 Prnd KO), 13 days (n = 15 Prnd WT, n = 14 Prnd KO), and 28 days (n = 14 Prnd WT, n = 14 Prnd KO) post-MCAO. While infarct volumes are similar at 3 days, Prnd KO mice exhibit significantly greater brain atrophy (loss of total brain volume) at 13 days and 28 days compared to WT controls, indicating progressive tissue loss during the chronic recovery phase. Data are presented as means ± SEM; *p < 0.05 using two-tailed Student’s t-test. **(C);** Functional neurological outcomes following stroke. Left panel: Neurological deficit scores assessed at 2 hours and 24 hours after MCAO show significantly worse deficits in Prnd KO mice compared to WT mice at 24 hours (n = 8 Prnd WT, n = 11 Prnd KO). Data are presented as means ± SEM; **p < 0.01 using one-way ANOVA with Tukey’s multiple-comparisons test. Right panel: Kaplan-Meier survival curve showing cumulative survival over 28 days post-MCAO. Prnd KO mice exhibit significantly reduced survival compared to WT controls (n = 25 Prnd WT, n = 40 Prnd KO). Statistical analysis by log-rank test.

Functional outcomes paralleled these anatomical findings. Neurological deficit scores were significantly elevated in Prnd KO mice compared to WT mice beginning at 24 hours post-stroke (Fig. 2C). Most critically, the survival rate of Prnd KO mice following ischemic stroke was substantially lower than that of WT mice (Fig. 2C). Collectively, these results indicate that while Prnd deficiency does not influence acute stroke injury, it significantly worsens long-term stroke outcomes, including tissue loss, neurological function, and survival.

### Prnd mRNA expression is upregulated in response to ischemic stroke

To characterize the temporal dynamics of Prnd expression following ischemic injury, we quantified Prnd mRNA levels in mouse brain tissue by quantitative reverse transcription PCR (qRT-PCR). In the ipsilateral (stroke-affected) hemisphere of Prnd WT mice, Prnd mRNA levels were significantly elevated at 3 days post-MCAO compared to baseline levels. This upregulation persisted through 13 days post-stroke but returned to basal levels by 28 days, suggesting a specific role for Prnd during the subacute recovery phase (Fig. 3A).

**Figure 3.**
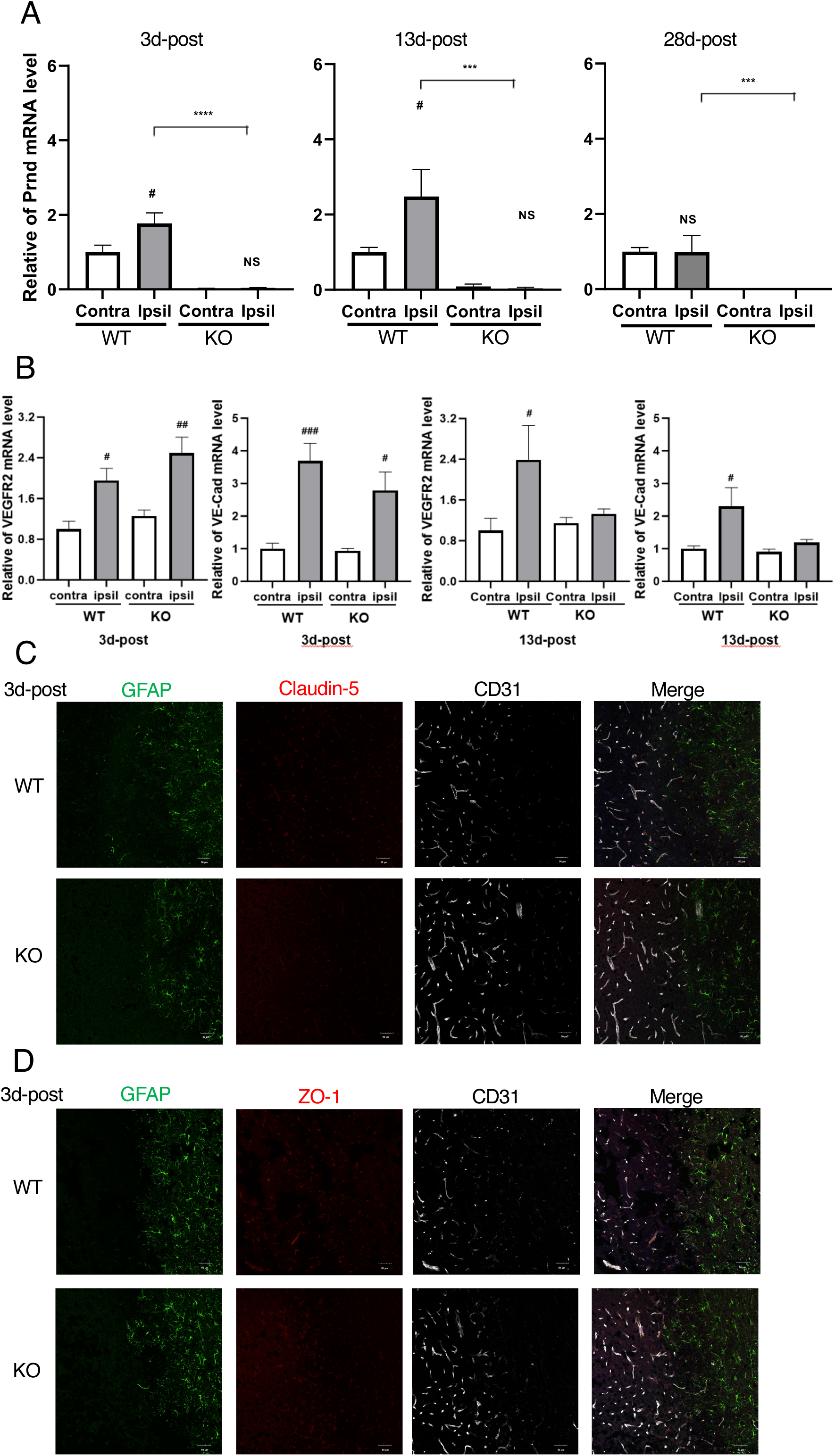
Prnd mRNA Expression Is Upregulated Following Stroke and Correlates with Pro-Angiogenic Gene Expression. **(A);** Temporal profile of Prnd mRNA expression in the mouse brain following ischemic stroke. Quantitative RT-PCR analysis of Prnd mRNA levels in ipsilateral (ischemic) and contralateral (non-ischemic) hemispheres at 3 days (n = 6 Prnd WT mice, n = 7 Prnd KO mice), 13 days (n = 7 Prnd WT mice, n = 8 Prnd KO mice), and 28 days(n = 7 Prnd WT mice, n = 8 Prnd KO mice) post-MCAO in Prnd WT mice. Prnd gene expression levels were normalized to GAPDH mRNA. The mean value of the contralateral hemisphere of Prnd WT mice was set as 1, and relative expression values were calculated for all other conditions. Prnd mRNA is significantly elevated in the ipsilateral hemisphere at 3 and 13 days post-stroke, returning to baseline by 28 days. Data are presented as means ± SEM; #p < 0.05, ##p < 0.01, ###p < 0.001 versus contralateral hemisphere of each genotype using one-way ANOVA with Tukey’s multiple-comparisons test. **(B);** Expression profiles of pro-angiogenic factors Kdr (encoding VEGFR2) and Cdh5 (encoding VE-Cadherin) mRNA in the ischemic brain at 3 days and 13 days post-MCAO. Gene expression levels were normalized to GAPDH, with the contralateral hemisphere of Prnd WT mice set as 1. Both Kdr/VEGFR2 and Cdh5/VE-Cadherin show increased expression in ipsilateral hemispheres of both genotypes at 3 days. However, sustained elevation is observed only in Prnd WT mice at 13 days, while expression returns to baseline in Prnd KO mice, suggesting Prnd is required for maintained pro-angiogenic signaling (n = 7 Prnd WT, n = 10 Prnd KO mice per time point). Data are presented as means ± SEM; #p < 0.05, ##p < 0.01, ###p < 0.001 versus contralateral hemisphere of each genotype using one-way ANOVA with Tukey’s multiple-comparisons test. **(C);** Representative immunofluorescence images showing claudin-5 expression (tight junction protein) in the ipsilateral cortex of Prnd WT and KO mice at 3 days post-MCAO. Claudin-5 expression is reduced in Prnd KO mice compared to WT controls, indicating compromised blood-brain barrier integrity. Scale bars, 50 μm. **(D);** Representative immunofluorescence images showing ZO-1 expression (tight junction scaffolding protein) in the ipsilateral cortex of Prnd WT and KO mice at 3 days post-MCAO. ZO-1 expression is markedly reduced in Prnd KO mice, further demonstrating impaired tight junction assembly. Scale bars, 50 μm.

To investigate how Prnd influences stroke-induced angiogenesis, we examined the expression of key pro-angiogenic factors in both Prnd WT and KO mice. We assessed mRNA levels of Kinase insert domain receptor (KDR) encoding vascular endothelial growth factor receptor 2 (VEGFR2) and Cdh5 (encoding vascular endothelial cadherin, VE-Cadherin). At 3 days post-MCAO, KDR/VEGFR2 mRNA levels were increased in the ipsilateral hemisphere of both Prnd WT and KO mice compared to the contralateral (unaffected) hemisphere.

Interestingly, KDR/VEGFR2 expression was even higher in Prnd KO mice than in WT mice at this early time point. However, the expression patterns diverged over time: while KDR/VEGFR2 mRNA levels remained elevated in the ipsilateral hemisphere of Prnd WT mice at 13 days post-MCAO, they returned to baseline levels in Prnd KO mice (Fig. 3B).

Similarly, Cdh5/VE-Cadherin mRNA levels increased in the ipsilateral hemisphere of both genotypes at 3 days post-MCAO. In Prnd WT mice, this elevation was sustained through 13 days post-stroke. In contrast, Cdh5/VE-Cadherin mRNA levels returned to baseline by 13 days in Prnd KO mice (Fig. 3B). These data suggest that Prnd expression is induced under hypoxic conditions characteristic of ischemic stroke, and that sustained Prnd expression is associated with prolonged elevation of pro-angiogenic factors including KDR/VEGFR2 and Cdh5/VE-Cadherin.

To assess Prnd-dependent BBB integrity defects at the molecular level, we examined the expression of key tight junction proteins. Claudin-5 is a transmembrane protein essential for maintaining BBB integrity [21], while zona occludens-1 (ZO-1) is a scaffolding protein crucial for assembling and stabilizing tight junction complexes [22]. We performed immunofluorescence staining for claudin-5 and ZO-1 in combination with the endothelial marker CD31 in coronal brain sections from Prnd WT and KO mice at 3 days post-MCAO. In the ischemic region, we observed moderately increased claudin-5 protein expression in Prnd WT mice compared to Prnd KO mice (Fig. 3C). More pronounced differences were observed for ZO-1, with substantially increased expression detected in the ischemic territory of Prnd WT mice relative to Prnd KO mice (Fig. 3D). These findings reveal that Prnd deficiency impairs BBB integrity following MCAO through impaired upregulation of critical endothelial tight junction proteins.

### Prnd deficiency worsens BBB breakdown and changes CNS blood vessel morphology after MCAO

To assess BBB disruption, we examined extravasation of mouse immunoglobulin G (IgG) into the brain parenchyma. IgG is a well-established marker of BBB breakdown following ischemic stroke [23]. In sham-operated control mice, IgG levels were comparable between Prnd WT and KO genotypes (Fig. 4A). However, at 3 days post-MCAO, IgG levels were significantly elevated in the ipsilateral striatum of Prnd KO mice compared to WT controls (Fig. 4B), confirming that Prnd deficiency exacerbates BBB disruption following ischemic injury.

**Figure 4.**
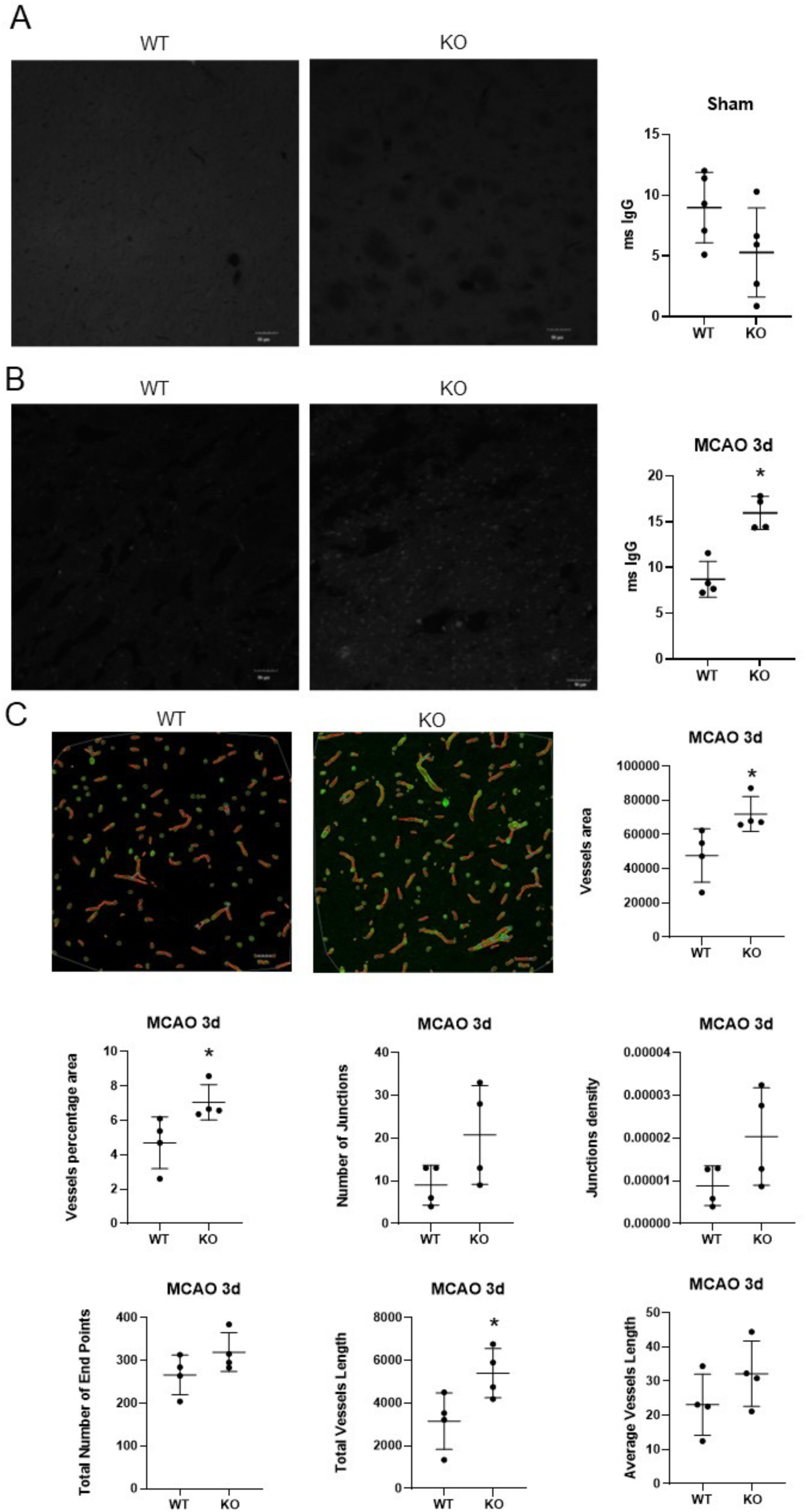
Prnd Deficiency Exacerbates Blood-Brain Barrier Disruption and Alters Cerebrovascular Morphology After Stroke. **(A);** Assessment of baseline BBB integrity in sham-operated (non-ischemic) Prnd WT and KO mice. Mouse immunoglobulin G (IgG) immunofluorescence intensity was quantified in brain parenchyma as a marker of BBB permeability. IgG levels are comparable between genotypes under basal conditions (n = 5 mice per group). Data are presented as means ± SEM using two-tailed Student’s t-test. **(B);** Quantification of BBB breakdown at 3 days post-MCAO. Mouse IgG extravasation into the ipsilateral striatum is significantly increased in Prnd KO mice compared to WT controls, indicating more severe BBB disruption in the absence of Prnd (n = 4 mice per group). Data are presented as means ± SEM; *p < 0.05 using two-tailed Student’s t-test. **(C);** Quantitative morphometric analysis of cerebrovascular networks in the ipsilateral cortex at 3 days post-MCAO. Parameters assessed include vessel area, vessel percentage area, and total vessel length. All measured parameters are significantly increased in Prnd KO mice compared to WT controls, suggesting aberrant vascular remodeling and disrupted vascular architecture in the absence of Prnd (n = 4 mice per group). Data are presented as means ± SEM; *p < 0.05 using two-tailed Student’s t-test.

To quantitatively assess acute vascular remodeling, we performed morphometric analysis of vascular networks in the ipsilateral cortex of Prnd WT and KO mice at 3 days post-MCAO using AngioTool. Several vascular parameters—including vessel area, vessel percentage area, and total vessel length- were increased in Prnd KO mice compared to WT mice (Fig. 4C).

In *Prnd* KO mice, the increase in vessel length and other parameters demonstrate chaotic sprouting mechanisms or greater stress during the acute phase of injury (3 days post-MCAO).

### Prnd deficiency worsens long-term neuroinflammation, neuron survival, and recovery of cerebral blood vessels after MCAO

To evaluate the long-term consequences of Prnd deficiency on neuroinflammation and neuronal survival, we performed immunofluorescence analysis of brain sections from Prnd WT and KO mice at 30 days post-MCAO. We used CD68 as a marker for activated macrophages/microglia and NeuN as a marker for mature neurons.

CD68 expression was markedly elevated in the ipsilateral hemisphere of Prnd KO mice compared to WT controls (Fig. 5A), indicating persistent neuroinflammation. Correspondingly, the number of NeuN-positive neurons was significantly reduced in the ipsilateral hemisphere of Prnd KO mice (Fig. 5A), indicative of decreased neuronal survival.

**Figure 5.**
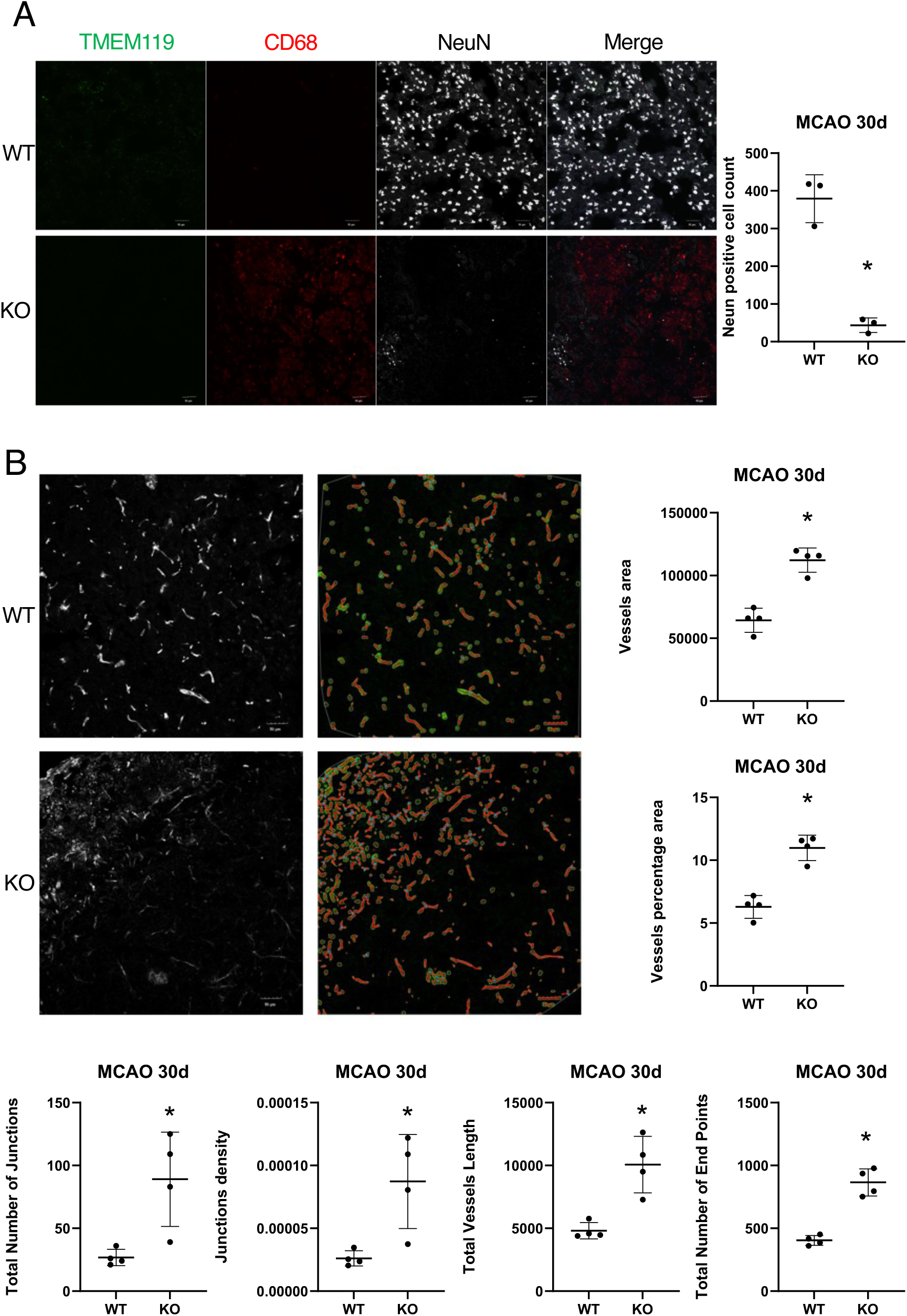
Prnd Deficiency Worsens Chronic Neuroinflammation, Reduces Neuronal Survival, and Impairs Cerebrovascular Recovery After Stroke. **(A);** Left panels: Representative immunofluorescence images showing TMEM119 (microglia marker), CD68 (macrophage/activated microglia marker) and NeuN (mature neuronal marker) staining in the ipsilateral cortex of Prnd WT and KO mice at 30 days post-MCAO. CD68 expression is elevated in Prnd KO mice, indicating persistent neuroinflammation. NeuN-positive neurons are reduced in Prnd KO mice, demonstrating decreased neuronal survival. Scale bars represent 50 μm. Right panel: Quantification of NeuN-positive cells in the ipsilateral cortex reveals significantly fewer surviving neurons in Prnd KO mice compared to WT controls at 30 days post-MCAO (n = 3 mice per group). Data are presented as means ± SEM; *p < 0.05 using two-tailed Student’s t-test. **(B);** Quantitative morphometric analysis of cerebrovascular networks in the ipsilateral cortex at 30 days post-MCAO. Parameters assessed include vessel area, vessel percentage area, total vessel length, and total number of junctions. All parameters are significantly increased in Prnd KO mice; however, qualitative assessment reveals severely disrupted vascular architecture compared to the relatively organized network in WT mice, indicating impaired functional vascular recovery (n = 4 mice per group). Data are presented as means ± SEM; *p < 0.05 using two-tailed Student’s t-test.

To assess long-term vascular recovery, we performed quantitative analysis of blood vessel networks in the ipsilateral cortex at 30 days post-MCAO using AngioTool. Multiple parameters were increased in Prnd KO mice compared to WT controls (Fig. 5B). However, qualitative examination revealed that the vascular network in Prnd KO mice exhibited disrupted architecture compared to the relatively normal vascular pattern in WT mice. These results demonstrate that Prnd deficiency impairs long-term cerebrovascular recovery following MCAO, resulting in abnormal blood vessel remodeling rather than functional restoration.

### Development of a Conditional Prnd Activation System Using Cre/lox Strategies

To achieve temporal and spatial control of Prnd expression in vivo, we generated a novel knock-in mouse model designated RD (Rosa26-Doppel). A targeting cassette was designed containing a CAG promoter (pCAG) followed by a loxP-flanked transcriptional STOP cassette (loxSTOPlox) upstream of the mouse Prnd cDNA coding sequence. This cassette was targeted to the ubiquitously expressed Rosa26 locus in mouse embryonic stem cells (Fig. 6A-C). In this configuration, Prnd remains transcriptionally repressed until Cre recombinase removes the STOP cassette, allowing Prnd expression under control of the CAG promoter. To validate this system in vitro, we infected cultured brain ECs from RD mice with adenovirus expressing Cre recombinase (Adenovirus-Cre). Cre infection resulted in activation of both Prnd mRNA (Fig. 6D) and protein (Fig. 6E) expression, confirming the functionality of the conditional allele. We also detected Prnd-dependent increases in expression of Hif1α and VEGF in cultured brain endothelial cells (Supp. Fig. 1).

**Figure 6.**
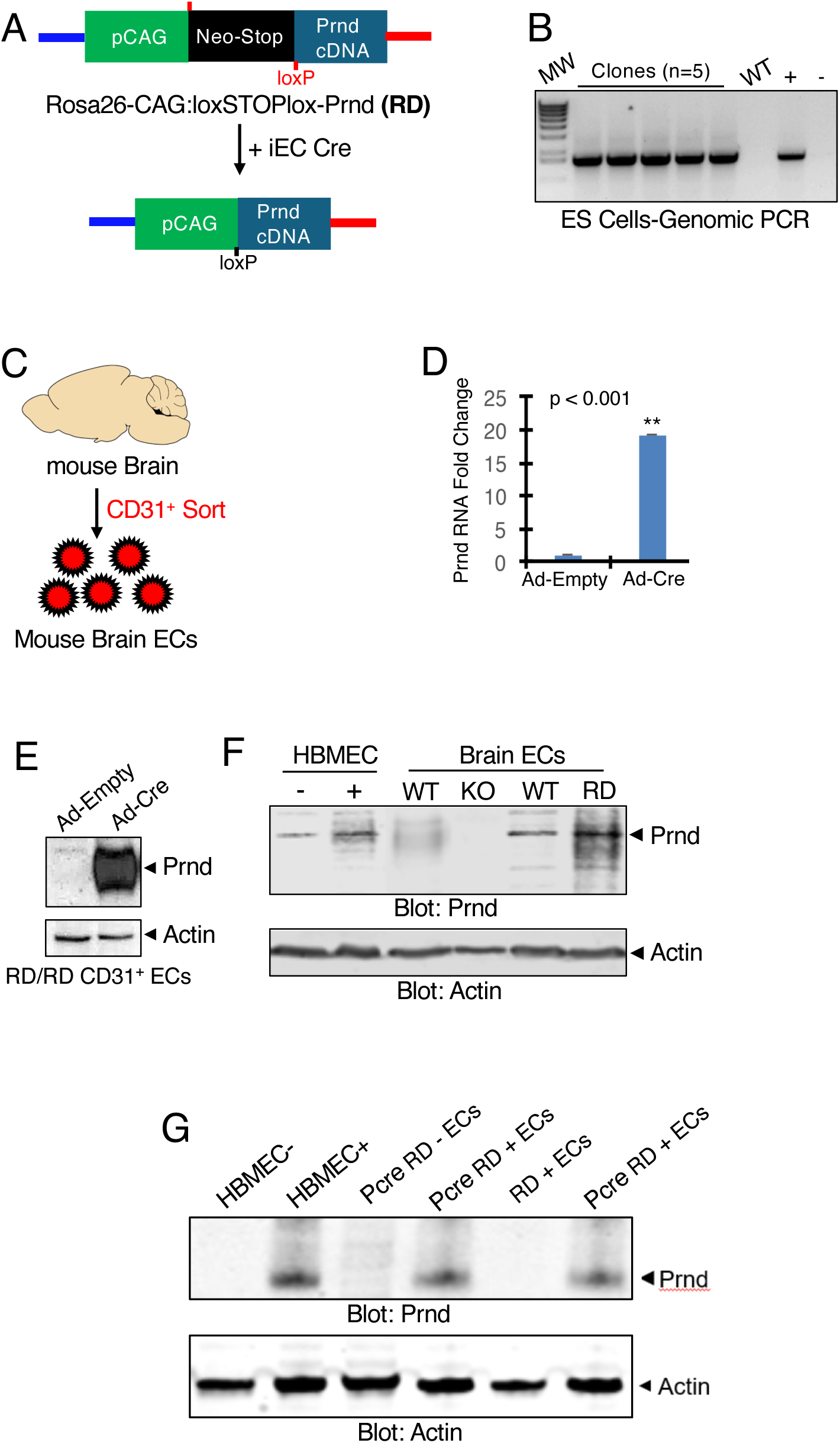
Development and Validation of a Conditional Prnd Expression System in Mouse Brain Endothelial Cells. **(A);** Schematic diagram of the Rosa26-Doppel (RD) targeting construct for generating a conditional Prnd knock-in mouse model. The targeting cassette contains a CAG promoter (pCAG) followed by a loxSTOPlox cassette positioned upstream of the mouse Prnd cDNA sequence. Upon Cre-mediated recombination, the STOP cassette is excised, allowing Prnd expression under control of pCAG. The construct was targeted to the ubiquitously expressed Rosa26 locus. **(B);** Genomic PCR validation of targeted embryonic stem (ES) cell clones. Genomic DNA was extracted from selected ES cell clones, and PCR was performed using primers F1 and R1 (positions indicated in A) to confirm proper integration of the targeting construct into the Rosa26 locus. **(C);** Schematic workflow showing isolation of brain endothelial cells (ECs) from adult RD/RD mice by fluorescence-activated cell sorting (FACS) using CD31 as an endothelial marker. **(D);** Quantitative RT-PCR analysis demonstrating activation of Prnd mRNA expression in cultured brain ECs from RD mice following infection with adenovirus expressing Cre recombinase (Adenovirus-Cre) compared to control-infected cells. Prnd expression was normalized to housekeeping gene. Data represent n = 3 independent experiments. **(E);** Western blot analysis confirming increased Doppel levels in cultured brain ECs from RD mice following Adenovirus-Cre infection, validating functional activation of the conditional allele at the protein level. **(F);** Comparative Western blot analysis of Doppel expression across three experimental systems: (i) human brain microvascular endothelial cells (HBMECs) infected with pLOC-PRND lentivirus versus control (left two lanes), demonstrating Prnd overexpression; (ii) CD31-positive primary brain ECs isolated from postnatal day 7 (P7) RD mice infected with AAV-Cre versus AAV-GFP control (middle two lanes); and (iii) CD31-positive ECs isolated from P12 RD/RD control mice versus PDGFB-CreERT2;RD/RD (Pcre-RD) mice that received tamoxifen injections at P1-P3 (right two lanes), confirming inducible endothelial-specific Prnd expression in vivo. **(G);** Western blot analysis validating sustained Prnd expression in adult mice. Comparison across: (i) HBMECs with lentiviral PRND overexpression (left two lanes); (ii) CD31-positive brain ECs from adult Pcre-RD mice with or without tamoxifen treatment at P1-P3 (middle two lanes); and (iii) CD31-positive ECs from adult RD/RD controls versus tamoxifen-treated Pcre-RD mice (right two lanes). Results confirm that tamoxifen-induced Cre activation during the early postnatal period leads to persistent Prnd expression in brain endothelial cells into adulthood.

For in vivo endothelial-specific activation, we intercrossed homozygous RD/RD mice with PDGFB-CreERT2 transgenic mice, in which tamoxifen-inducible Cre recombinase is expressed specifically in vascular endothelial cells [24, 25]. The resulting PDGFB-CreERT2/+;RD/RD mice (designated Pcre-RD) or RD/RD control littermates (Cre-negative) were injected intragastrically with tamoxifen from P1 to P3, with analysis performed at P12. CD31-positive endothelial cells isolated from Pcre-RD brains showed elevated Doppel expression compared to RD/RD control mice that lacked the Cre transgene (Fig. 6F).

To confirm endothelial-specific and sustained Prnd expression into adulthood, we analyzed adult Pcre-RD mice that had received tamoxifen treatment at P1-P3. CD31-positive endothelial cells from adult Pcre-RD brains exhibited increased Doppel expression compared to both RD/RD controls and Pcre-RD mice that had not received tamoxifen (Fig. 6G). These results confirm that the PDGFB-CreERT2;RD system enables tamoxifen- inducible activation of Prnd expression in brain vascular endothelial cells.

To determine whether endothelial-specific activation of Prnd influences acute stroke injury, we measured infarct volumes in RD control and Pcre-RD mice at 24 hours following MCAO using 2,3,5-triphenyltetrazolium chloride (TTC) staining of coronal brain sections. Infarct volumes were comparable between the two groups (Fig. 7A), indicating that endothelial Prnd activation does not significantly alter acute ischemic damage. To characterize the temporal pattern of Doppel expression following stroke, we performed immunofluorescence analysis at multiple time points in RD and Pcre-RD mice. While Prnd expression showed minimal changes at early time points, we detected an increase in Doppel expression in Pcre-RD mice at 7 days post-MCAO (Fig. 7B). This peak in Prnd expression was accompanied by a concurrent elevation in vascular endothelial growth factor (VEGF) and laminin protein levels (Fig. 7B-D). These findings demonstrate that endothelial activation of Prnd induces a temporally regulated peak in Doppel expression during the subacute recovery phase, which in turn promotes expression of the critical pro-angiogenic factor VEGF and BBB integrity protein laminin. Laminin is an essential extracellular matrix (ECM) protein that is predominantly localized to the vascular basement membrane, where it contributes to structural and functional integrity [26]. Specificity of the anti-Prnd antibody was confirmed using KO brain tissue and tamoxifen-treated Pcre-RD brain tissue not exposed to MCAO (Supp. Fig. 2).

**Figure 7.**
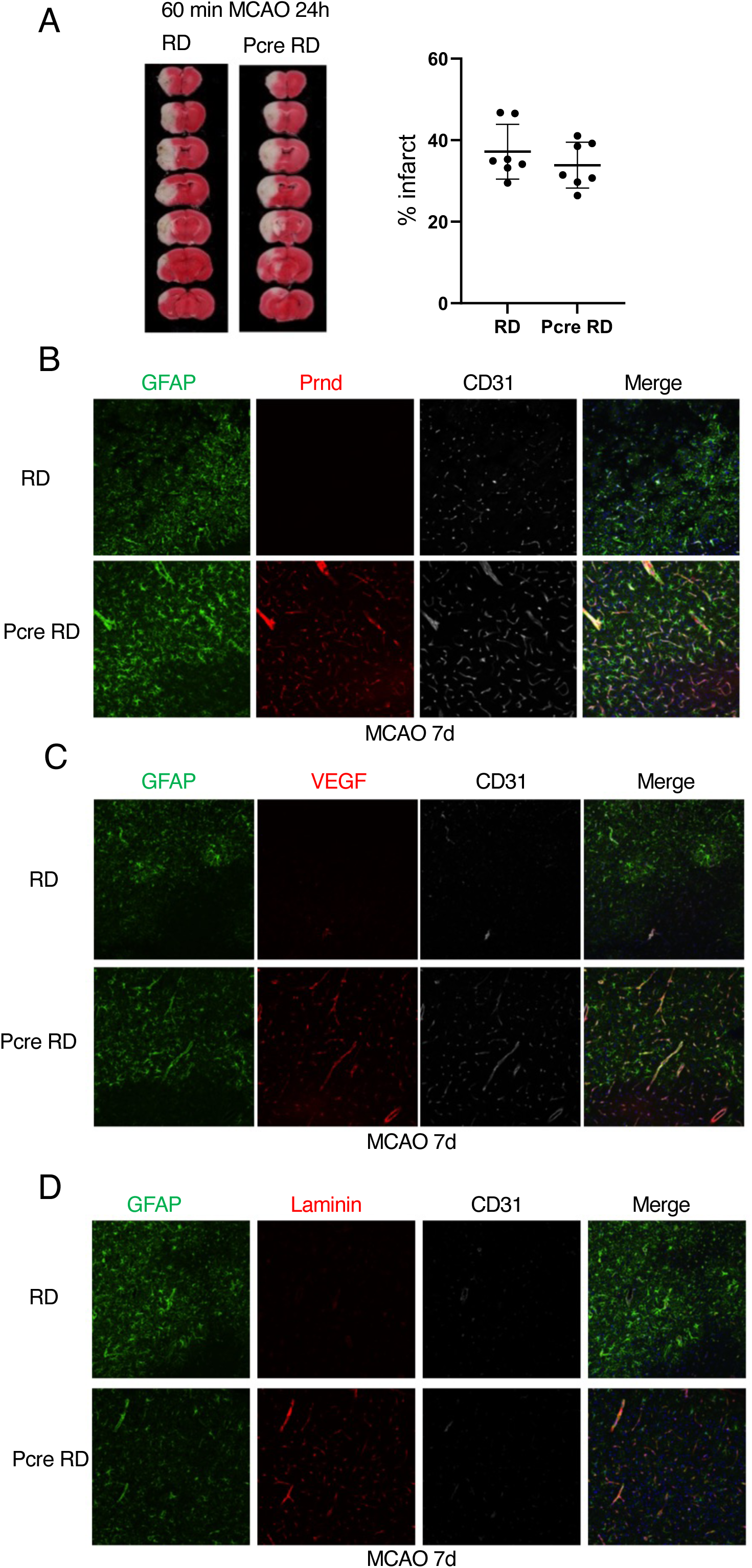
Endothelial Prnd Activation Does Not Affect Acute Stroke Injury but Induces Temporally Regulated Angiogenic Gene Expression. **(A);** Assessment of acute brain injury in RD control and Pcre-RD mice at 24 hours following 60 minutes of MCAO. Infarct volumes were quantified in TTC-stained 1 mm coronal brain sections. Infarct volumes are comparable between genotypes, indicating that endothelial-specific Prnd activation does not significantly alter acute ischemic damage (n = 7 mice per group). Data are presented as means ± SEM using two-tailed Student’s t-test. **(B);** Representative immunofluorescence images showing GFAP (astrocyte marker) and Prnd expression in the ipsilateral cortex of RD control and Pcre-RD mice at 7 days post-MCAO. An increase in Doppel expression is observed in Pcre-RD mice, demonstrating temporally regulated Prnd activation during the subacute recovery phase (n = 3 mice per group). Scale bars represent 50 μm. **(C);** Representative immunofluorescence images showing GFAP and vascular endothelial growth factor (VEGF) expression in the ipsilateral cortex of RD and Pcre-RD mice at 7 days post-MCAO. The peak in Prnd expression (B) is accompanied by a concurrent elevation in VEGF protein levels in Pcre-RD mice, indicating that Prnd activation promotes expression of this critical pro-angiogenic factor (n = 3 mice per group). Scale bars represent 50 μm. **(D);** Representative immunofluorescence images showing GFAP and laminin expression in the ipsilateral cortex of RD and Pcre-RD mice at 7 days post-MCAO. The peak in Prnd expression (B) is accompanied by a concurrent elevation in laminin protein levels in Pcre-RD mice, indicating that Prnd activation promotes expression of this crucial BBB protein (n = 3 mice per group). Scale bars represent 50 μm.

### Forced expression of Doppel promotes BBB recovery

To investigate whether elevated Prnd expression promotes BBB recovery, we examined the expression of key BBB-associated proteins in the ipsilateral striatum at 7 days post-MCAO. ZO-1 and claudin-5 are critical tight junction proteins that collaborate in brain endothelial cells to establish and maintain BBB integrity [27, 28]. We performed comprehensive immunofluorescence analysis of Prnd, ZO-1, and claudin-5 in brain sections from Pcre-RD and RD control mice at 7 days post-MCAO. As expected, Doppel expression was markedly elevated in Pcre-RD mice (Fig. 8A). Importantly, this increase in Prnd was associated with significantly enhanced expression ZO-1 (Fig. 8B), and claudin-5 (Fig. 8C) in the ipsilateral striatum of Pcre-RD mice compared to RD controls. These results demonstrate that peak Prnd expression induces coordinate upregulation of proteins essential for BBB integrity, suggesting that Prnd promotes BBB recovery during the subacute phase following ischemic injury. We did not detect differences in blood vessel morphologies in tamoxifen-treated Pcre-RD mouse brains exposed to sham treatment (Supp. Fig. 3).

**Figure 8.**
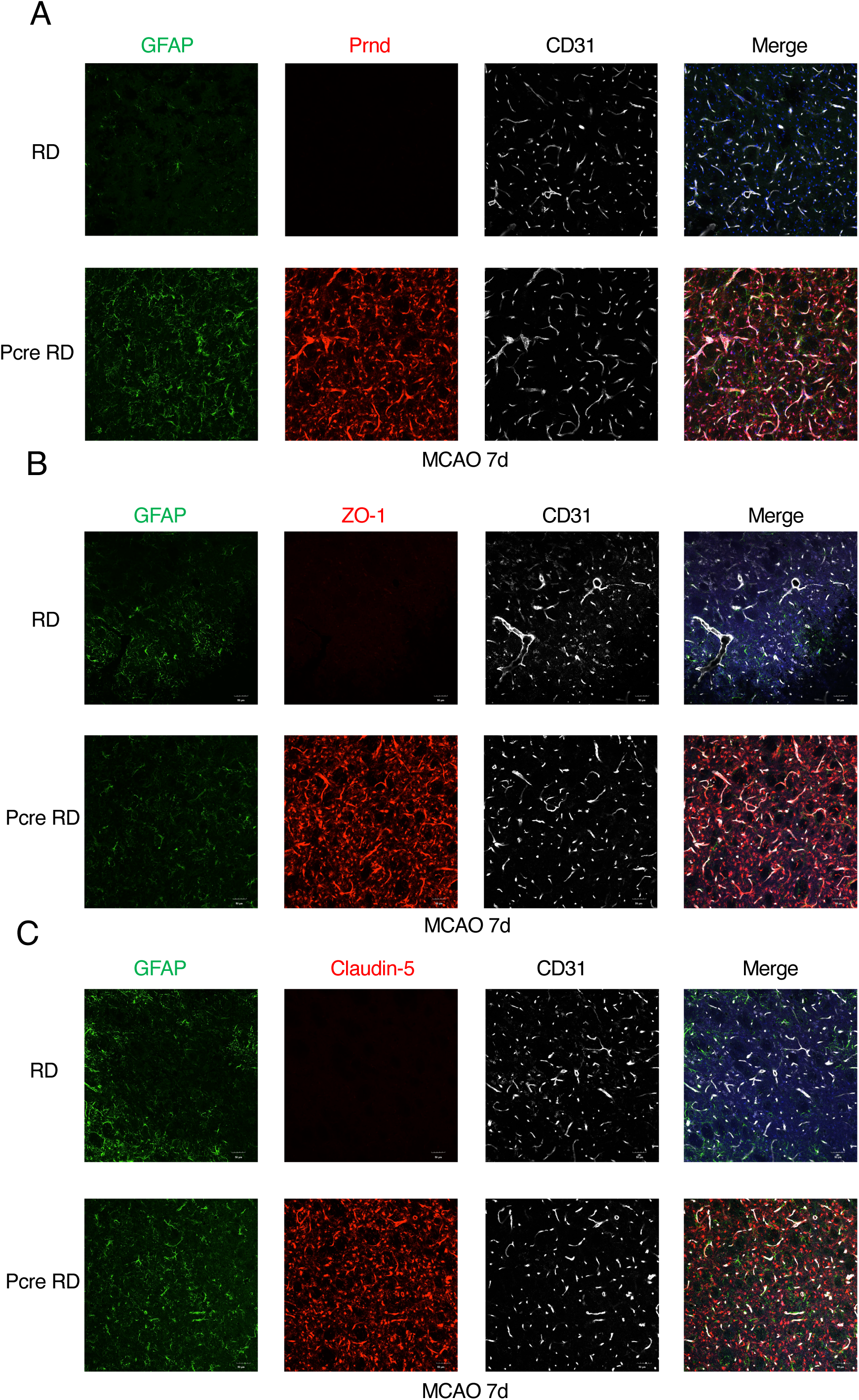
Forced Endothelial Prnd Expression Promotes Blood-Brain Barrier Recovery Following Ischemic Stroke. **(A);** Representative immunofluorescence images of Doppel expression in the ipsilateral striatum of RD control and Pcre-RD mice at 7 days post-MCAO. Prnd expression is markedly elevated in Pcre-RD mice, confirming successful activation of the conditional allele in the post-stroke brain (n = 3 mice per group). Scale bars, 50 μm. **(B);** Representative immunofluorescence images of zona occludens-1 (ZO-1) expression in the ipsilateral striatum at 7 days post-MCAO. ZO-1, a critical tight junction scaffolding protein, is substantially upregulated in Pcre-RD mice, demonstrating improved tight junction assembly (n = 3 mice per group). Scale bars, 50 μm. **(C);** Representative immunofluorescence images of claudin-5 expression in the ipsilateral striatum at 7 days post-MCAO. Claudin-5, a transmembrane tight junction protein essential for BBB integrity, shows enhanced expression in Pcre-RD mice. Together with B and C, these results demonstrate that Prnd activation promotes coordinate upregulation of key BBB-associated proteins, facilitating BBB recovery (n = 3 mice per group). Scale bars, 50 μm.

To investigate the role of Prnd in cerebrovascular morphology recovery after ischemic stroke, we used AngioTool to perform quantitative morphometric analysis of brain vascular networks at 7 days post MCAO in RD control and Pcre-RD mice ipsilateral cortex using anti-CD31 immunofluorescence imaging. We assessed multiple vascular parameters including vessel area, percentage of vessels per area, total vessel length, and average vessel length. Our analysis revealed significant increases in almost all parameters in Pcre-RD mice compared to RD controls (Fig. 9A, B). These findings demonstrate that Prnd promotes cerebrovascular morphology recovery after ischemic stroke.

**Figure 9.**
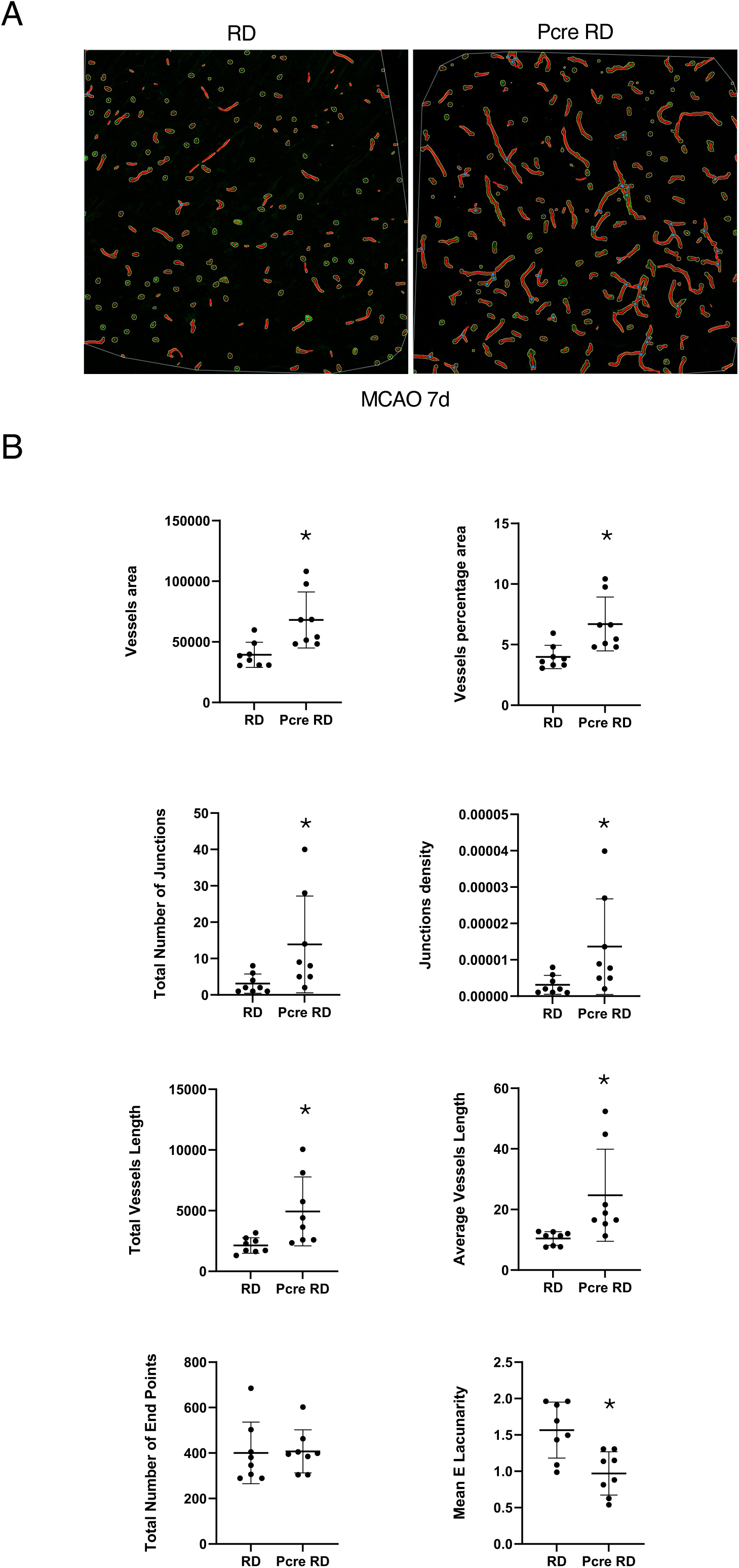
Endothelial Prnd Expression Enhances Post-Stroke Angiogenesis. **(A);** Representative immunofluorescence images from coronal brain sections through the ipsilateral cortex of RD and Pcre-RD mice at 7 days post-MCAO, stained with anti-CD31 antibodies. Enhanced vascular network density is evident in Pcre-RD mice during the active angiogenic phase following stroke. Scale bars represent 50 μm. **(B);** Quantitative morphometric analysis of vascular networks in the ipsilateral cortex at 7 days post-MCAO. Parameters include vessel area, vessel percentage area, total vessel length, average vessel length, and number of junctions. Multiple vascular parameters are significantly increased in Pcre-RD mice, demonstrating that Prnd activation enhances post-stroke angiogenesis and vascular remodeling (n = 8 mice per group). Data are presented as means ± SEM; *p < 0.05 using two-tailed Student’s t-test.

### Activation of Prnd promotes long-term neuron survival after ischemic stroke

To assess whether endothelial Prnd activation influences long-term neuronal outcomes, we quantified neuronal survival in the ipsilateral cortex of Pcre-RD and RD mice at 14 and 30 days post-MCAO. Immunofluorescence staining was performed using ms IgG, TMEM119 (a specific marker for resident microglia), CD68 (a marker for activated macrophages/microglia), and NeuN (a marker for mature neurons), followed by quantification of NeuN-positive cells. At 14 days post-MCAO, the ipsilateral cortex of Pcre-RD mice contained significantly more NeuN-positive neurons compared to RD controls (Fig. 10A, B), indicating improved neuronal survival. At 30 days post-MCAO, Pcre-RD mice continued to show a trend toward increased numbers of NeuN-positive neurons in the ipsilateral cortex compared to RD mice, although this difference did not reach statistical significance (Fig. 10C, D). These findings reveal that endothelial-specific activation of Prnd promotes neuronal survival during the chronic recovery phase following ischemic stroke, likely through improved vascular function and BBB integrity.

**Figure 10.**
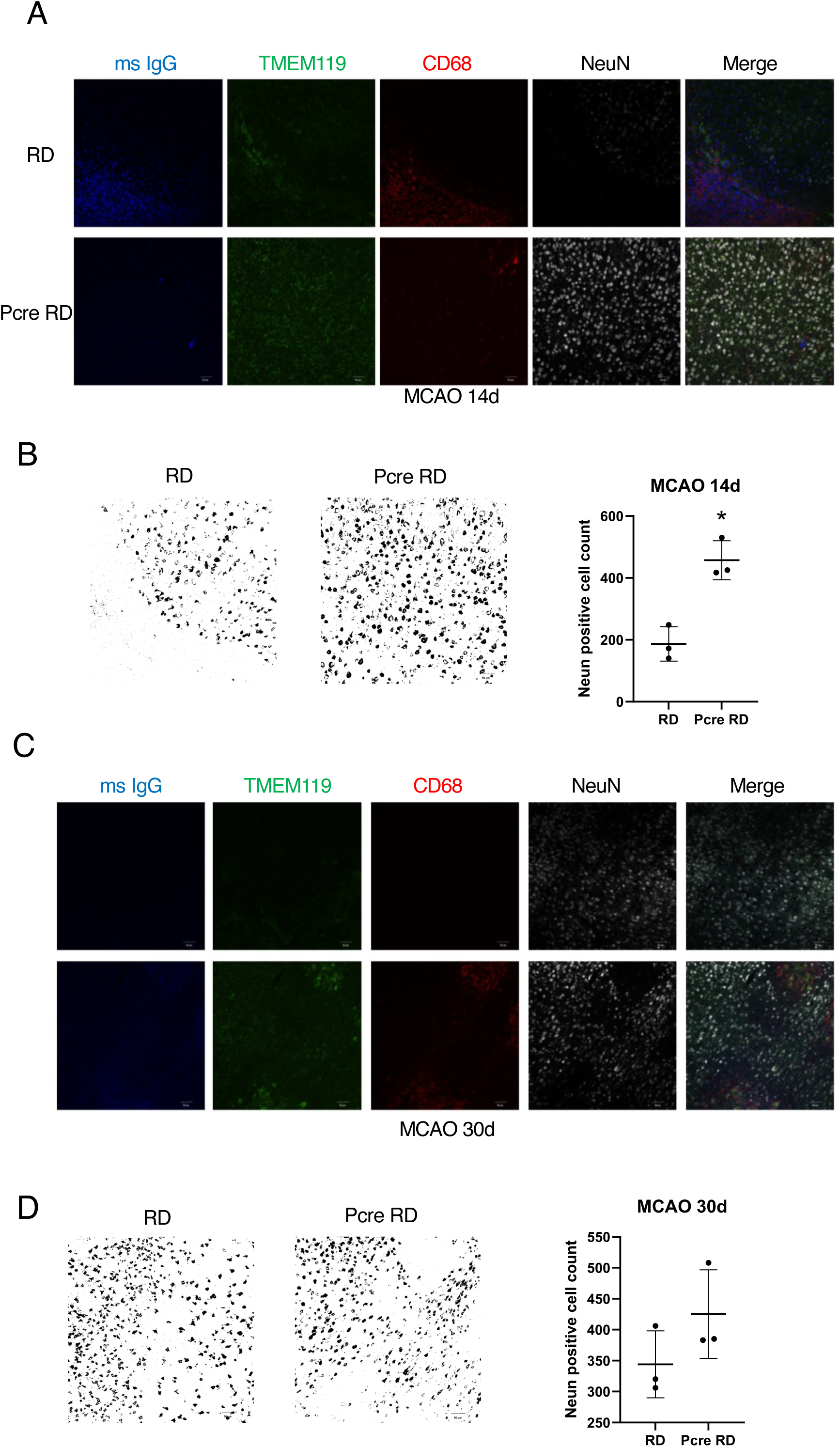
Endothelial Prnd Expression Promotes Long-Term Neuronal Survival Following Ischemic Stroke. **(A);** Representative immunofluorescence images showing ms IgG, TMEM119 (resident microglia marker), CD68 (macrophage/activated microglia marker), and NeuN (mature neuronal marker) expression in the ipsilateral cortex of RD control and Pcre-RD mice at 14 days post-MCAO. Enhanced NeuN staining in Pcre-RD mice indicates improved neuronal survival during the subacute recovery phase (n = 3 mice per group). Scale bars represent 100 μm. **(B);** Quantification of NeuN-positive neurons in the ipsilateral cortex at 14 days post-MCAO. Pcre-RD mice exhibit significantly more surviving neurons compared to RD controls, demonstrating that endothelial Prnd activation confers neuroprotection during the recovery phase (n = 3 mice per group). Data are presented as means ± SEM; *p < 0.05 using two-tailed Student’s t-test. **(C);** Representative immunofluorescence images showing ms IgG, TMEM119, CD68, and NeuN expression in the ipsilateral cortex of RD and Pcre-RD mice at 30 days post-MCAO, during the chronic recovery phase (n = 3 mice per group). Scale bars represent 50 μm. **(D);** Quantification of NeuN-positive neurons in the ipsilateral cortex at 30 days post-MCAO. Pcre-RD mice show a trend toward increased neuronal survival compared to RD controls, though the difference does not reach statistical significance at this later time point (n = 3 mice per group). Data are presented as means ± SEM using two-tailed Student’s t-test. These results suggest that the neuroprotective benefits of Prnd activation are most pronounced during the subacute recovery phase.

Lastly, to investigate the pro-angiogenic effects of Prnd on endothelial cells, we performed gain-of-function experiments in two complementary systems: human brain microvascular endothelial cells (HBMECs) and primary brain endothelial cells isolated from postnatal day 7 (P7) RD mice. HBMECs were infected with lentivirus expressing human PRND (pLOC-PRND) or with control lentivirus expressing GFP/RFP (pLOC negative control). Cells were harvested 48 hours post-infection for Western blot analysis. P7 RD mouse brain endothelial cells were infected with either AAV-Cre (to activate endogenous Prnd) or AAV-GFP (control). Cells were harvested 72 hours post-infection for immunofluorescence analysis. In RD mouse brain endothelial cells, AAV-Cre infection resulted in increased expression of Prnd, ZO-1 and VE-Cadherin compared to AAV-GFP-infected controls (Supp. Fig. 1A and Supp. Fig. 4A, 4B). In HBMECs, Prnd overexpression led to increased intracellular VEGF, p-VEGF R2 levels and decreased intracellular angiostatin (an endogenous angiogenesis inhibitor) (Supp. Fig. 4C). Furthermore, Prnd overexpression resulted in enhanced phosphorylation of multiple signaling molecules that promote angiogenesis, including focal adhesion kinase (FAK), protein kinase B (Akt), extracellular signal-regulated kinases 1 and 2 (Erk1/2), and SRC proto-oncogene kinase (SRC) (Supp. Fig. 4C). Collectively, these results demonstrate that Prnd activation or overexpression promotes pro-angiogenic signaling pathways in endothelial cells in vitro, providing mechanistic insight into its beneficial effects on cerebrovascular recovery following ischemic stroke.

## Discussion

Post-stroke neovascularization is temporally regulated process that initiates within hours of ischemic injury through the activation of EC proliferation and sprouting mechanisms driven by local cues within the ischemic penumbra [29, 30]. The emergence of newly formed blood vessels is evident in peri-infarct regions by four days post-stroke, with angiogenesis peaking between 7 and 21 days and persisting for weeks to months, thus contributing to tissue repair and functional neurological recovery [31, 32]. The mechanistic understanding of post-ischemic angiogenesis remains incomplete, with accumulating evidence suggesting that angiogenesis serves a temporally biphasic role in stroke pathophysiology. During the acute phase (defined as the first 72 hours post-ischemia), the upregulation of growth factors including VEGF and MMPs promotes increased vascular permeability through disruption of inter-endothelial junctional complexes [33]. This compromise of BBB integrity exacerbates vasogenic edema formation and elevates the risk of hemorrhagic transformation [33]. Conversely, during the subacute and chronic phases, angiogenesis transitions to a predominantly regenerative process that is critical for long-term neurological recovery [34]. These newly established vascular networks enhance regional hemodynamics, facilitate oxygen and nutrient delivery to metabolically compromised tissue, and promote the reconstruction of functional NVUs.

The present investigation demonstrates that neither Prnd deficiency (loss-of-function) nor constitutive Prnd activation (overexpression) significantly altered short-term stroke outcomes, suggesting that Prnd does not play a critical role in the acute pathophysiological cascades. However, our findings reveal that Prnd deficiency markedly impaired long-term stroke outcomes through mechanisms involving compromised NVU recovery and diminished neuronal survival in the post-acute period following MCAO. Molecular analysis revealed that Prnd mRNA expression is dynamically upregulated in response to ischemic stroke, with sustained elevation of Prnd transcript levels correlating temporally with increased expression of pro-angiogenic molecular markers including Kdr/VEGFR2 and Cdh5/VE-cadherin. These findings suggest coordinated transcriptional regulation of Prnd-dependent angiogenic programs. Notably, Prnd deficiency substantially attenuated the upregulation of pro-angiogenic factors in the ipsilateral hemisphere at 13 days post-MCAO. Collectively, these data reveal that Prnd plays a necessary role in sustaining the angiogenic response during the critical proliferative phase of post-stroke blood vessel remodeling.

Gain-of-function experiments utilizing overexpression of Prnd at 7 days post-MCAO, corresponding to the peak angiogenic window [29] revealed coordinated upregulation of multiple proteins essential for angiogenesis and vascular integrity, including VEGF, the basement membrane component laminin, and the tight junction proteins ZO-1 and claudin-5. VEGF stimulates neovascularization during embryonic development and in pathological conditions including tumor growth [35]. The biological effects of VEGF are mediated through binding to cognate receptor tyrosine kinases on ECs, triggering intracellular signaling cascades that promote ECs survival and vascular permeability [36]. The concurrent upregulation of tight junction proteins ZO-1 and claudin-5 alongside VEGF following Prnd activation suggests a mechanism whereby Prnd may promote angiogenesis while simultaneously facilitating BBB restoration, potentially mitigating increased vascular permeability associated with acute-phase VEGF expression. Functional studies confirmed that Prnd activation promoted angiogenesis during the proliferative phase and enhanced long-term neuronal survival following MCAO, supporting a neuroprotective role for Prnd-mediated vascular regeneration. Collectively, these data substantially expand our mechanistic understanding of Prnd functions in ischemic stroke pathophysiology and position Prnd/Doppel as a potential therapeutic target for modulating angiogenesis during the proliferative phase and promoting long-term recovery in cerebral ischemia.

Our previous investigations revealed that Prnd/Doppel expression is abundant in angiogenic blood vessels within the developing mouse brain but becomes virtually undetectable in quiescent vasculature of the adult brain [12], suggesting specific developmental regulation of Prnd expression. Consistent with these prior observations, the current study detected negligible Doppel expression in adult wild-type and Prnd-/- mice under physiological conditions, and constitutive activation of the Prnd gene in sham-operated adult Pcre-RD mice produced only modest increases in Doppel within brain ECs. We hypothesize that in the absence of active angiogenic signaling, Doppel exhibits inherent structural instability and undergoes rapid degradation through cellular proteolytic machinery. This post-translational regulation would account for the paradoxically low Doppel levels despite documented increases in mRNA transcripts under non-angiogenic conditions. Supporting the EC-specific expression of Prnd, our RT-PCR analyses confirmed that Prnd transcripts were exclusively detected in CD31-positive cells isolated from brain tissue. Furthermore, genetic activation of Prnd enhanced both mRNA and protein levels in developing mouse brain ECs as well as in cultured EC lines, confirming the capacity for Prnd expression under appropriate cellular contexts. The temporal dynamics of Doppel expression following MCAO further support the hypothesis that Prnd participates specifically in active angiogenic processes. We observed substantial Prnd accumulation exclusively at 7 days post-MCAO in Pcre-RD mice. This expression pattern aligns with the established timeline of peak angiogenic activity following ischemic stroke, suggesting that Prnd expression is tightly coupled to active vessel formation and may be dispensable once vascular remodeling is complete.

While our results demonstrate that Prnd expression promotes angiogenesis during the proliferative phase and enhances long-term neuronal survival following MCAO, several important questions warrant further investigation. First, it remains to be determined whether Prnd activation can improve long-term survival rates and facilitate cognitive rehabilitation following ischemic stroke, outcomes of critical clinical relevance. Given that Prnd enhances both vascular regeneration and neuronal survival, it is plausible that Prnd-mediated improvements in tissue repair could translate to better functional recovery and reduced long-term disability.

Longitudinal behavioral studies incorporating comprehensive neurocognitive assessments would be valuable for evaluating the full therapeutic potential of Prnd modulation in stroke recovery. Second, although CD31 serves as a reliable marker for brain ECs, CD31 expression is not exclusively restricted to the endothelial lineage. This glycoprotein is also expressed by platelets and various leukocyte populations, including monocytes, neutrophils, and lymphocyte subsets [37, 38], all of which infiltrate ischemic brain tissue as part of the post-stroke inflammatory response. Consequently, we cannot definitively exclude the possibility that these non-endothelial CD31-positive cells may also express Prnd and contribute to the observed effects. Future investigations employing more refined cell-type-specific approaches, such as single-cell RNA sequencing or immunofluorescence co-localization studies with multiple EC-specific markers, will be essential for unambiguously identifying the cellular sources of Prnd expression. Additionally, comprehensive transcriptomic and proteomic profiling of ECs following Prnd activation in the post-ischemic brain would provide valuable insights into the downstream molecular pathways through which Prnd exerts its pro-angiogenic effects, potentially revealing additional therapeutic targets within the Prnd-regulated angiogenic network.

## Acknowledgements and Funding Statement

Research reported in this manuscript was supported, in part, by the National Institute of Neurological Disease and Stroke of the National Institutes of Health (R01NS087635 and R01NS122143). The authors acknowledge the use of resources and technical support from CCSG-funded core facilities as well as the Cancer Neuroscience Program at MD Anderson Cancer Center. The content in this manuscript is solely the responsibility of the authors and does not necessarily represent the official views of the National Institutes of Health.

---

**Supplemental Figure 1.**
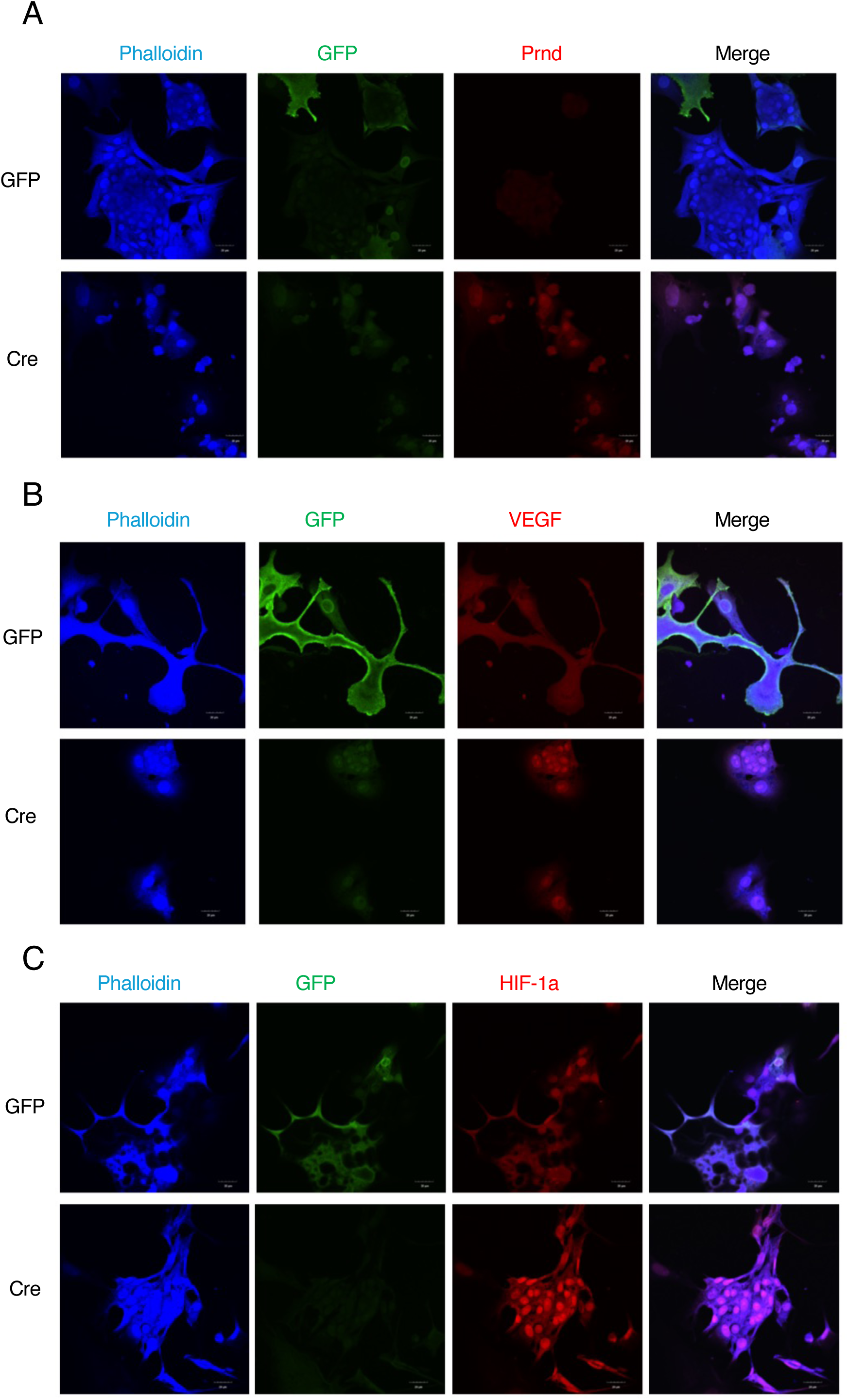
Prnd Activation Enhances Expression of Pro-Angiogenic Factors In Vitro. **(A);** Representative immunofluorescence images showing Prnd expression in cultured brain endothelial cells isolated from P7 RD mice, and infected with AAV-Cre or AAV-GFP control. Prnd expression is observed following AAV-Cre infection, validating efficient Cre-mediated recombination in vitro (n = 3 independent experiments). Scale bars, 50 μm. **(B);** Representative immunofluorescence images showing vascular endothelial growth factor (VEGF) expression in P7 RD brain endothelial cells infected with AAV-Cre versus AAV-GFP. Prnd activation leads to increased VEGF expression, demonstrating a direct pro-angiogenic effect at the cellular level (n = 3 independent experiments). Scale bars, 50 μm. **(C);** Representative immunofluorescence images showing hypoxia-inducible factor 1-alpha (HIF-1α) expression in P7 RD brain endothelial cells infected with AAV-Cre versus AAV-GFP. Enhanced HIF-1α expression following Prnd activation suggests engagement of hypoxia-responsive pro-angiogenic pathways (n = 3 independent experiments). Scale bars, 50 μm.

**Supplemental Figure 2.**
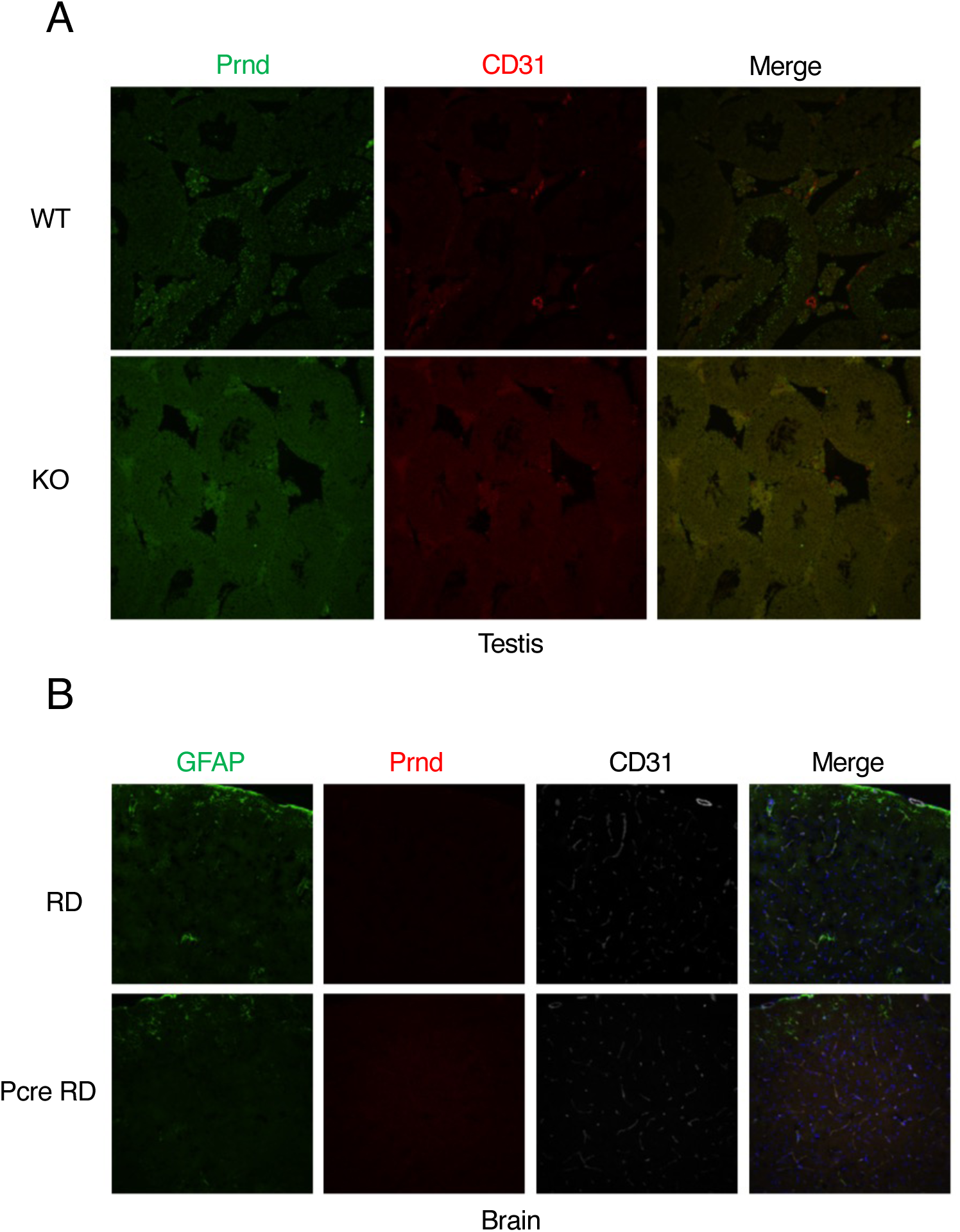
Analysis of Doppel Expression in Mouse Testis and Brain. **(A);** Representative immunofluorescence images showing Prnd and CD31 (endothelial marker) expression in the testis of Prnd WT and KO mice. Images demonstrate Prnd expression in WT but not in KO testis. Scale bars represent 50 μm. **(B)**; Representative immunofluorescence images showing GFAP, Prnd, and CD31 expression in RD control and Pcre RD mice brain. Increased Prnd expression in Pcre RD mice brain indicates activation of Prnd at P3-P5 promotes Prnd expression (n = 3 mice per group). Scale bars represent 50 μm.

**Supplemental Figure 3.**
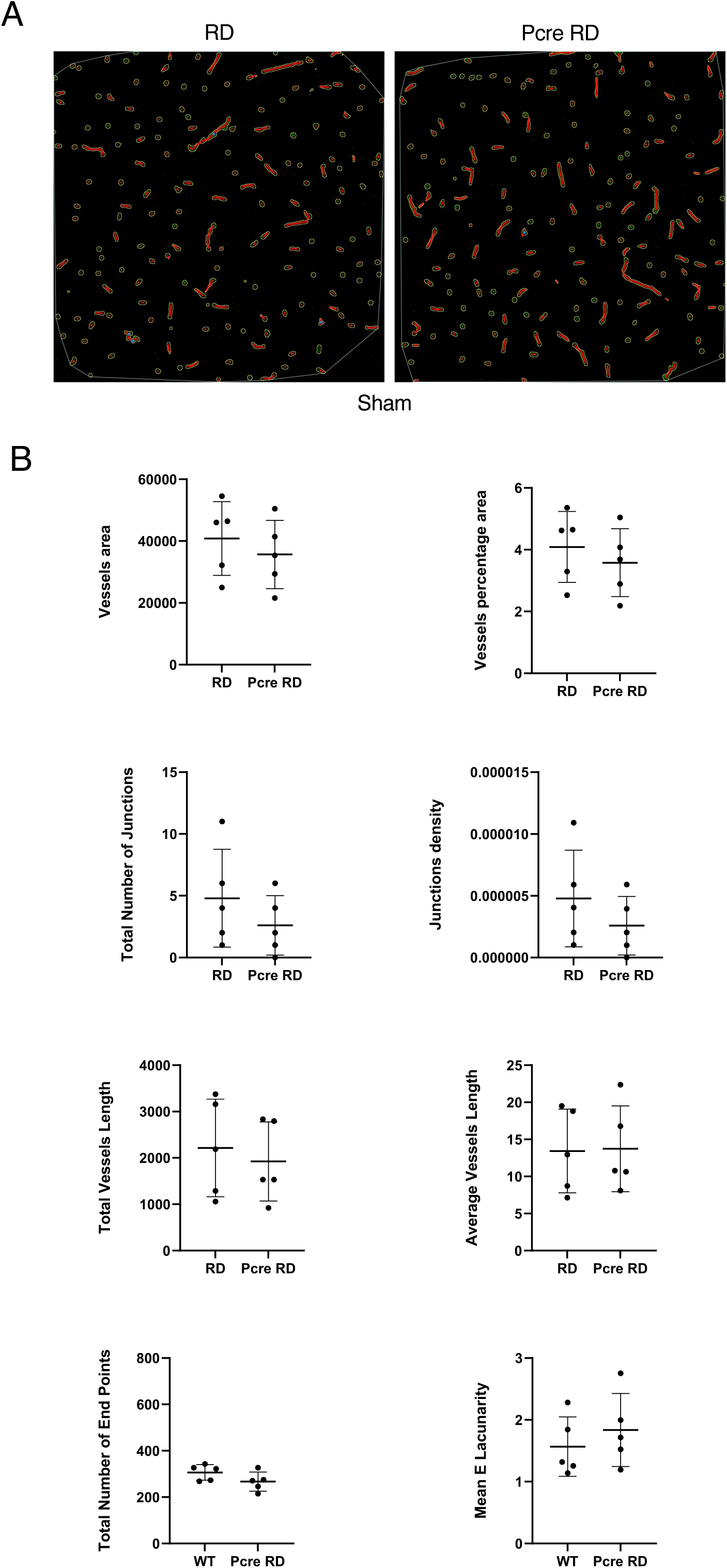
Endothelial Prnd Activation has no Effect on Baseline Cerebrovascular Development. **(A);** Representative immunofluorescence images from coronal brain sections through the striatum of 12-week-old RD control and Pcre-RD mice (tamoxifen-treated at P1-P3). Sections were stained with anti-CD31 antibodies to visualize the vascular network. Images demonstrate similar vascular density in Pcre-RD mice under baseline (non-ischemic) conditions. Scale bars represent 50 μm. **(B);** Quantitative morphometric analysis of vascular networks in the striatum of 12-week-old RD and Pcre-RD mice. Parameters assessed include vessel area, vessel percentage per area, total vessel length, average vessel length, and number of junctions. Multiple vascular parameters are similar between RD control and Pcre-RD mice, indicating that early postnatal endothelial Prnd activation has no effect on cerebrovascular development and maturation under physiological conditions (n = 5 mice per group). Data are presented as means ± SEM; *p < 0.05 using two-tailed Student’s t-test.

**Supplemental Figure 4.**
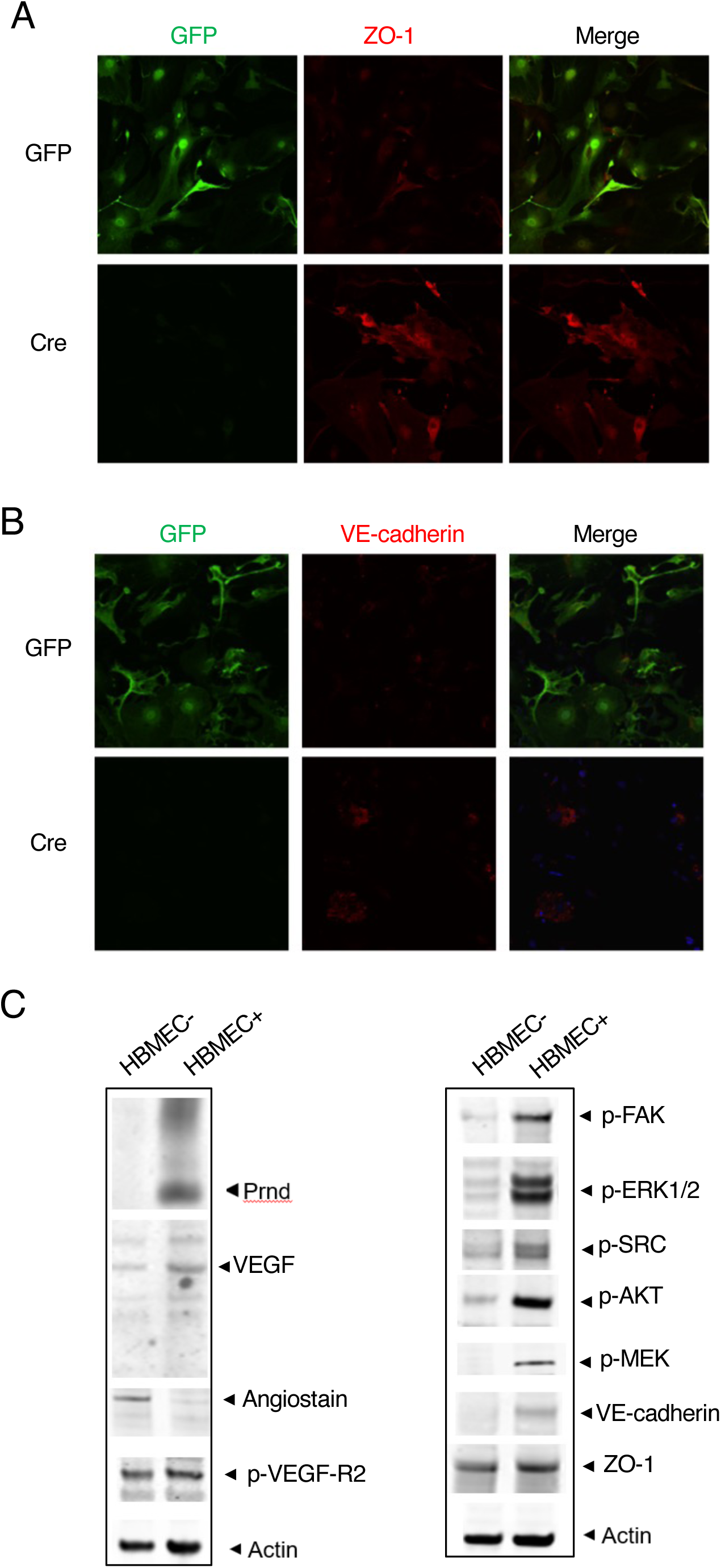
Prnd Overexpression Promotes Pro-Angiogenic Signaling in Endothelial Cells In Vitro. **(A);** Representative immunofluorescence images showing VE-Cadherin (vascular endothelial cadherin) expression in P7 RD mouse brain endothelial cells infected with AAV-Cre versus AAV-GFP control. Prnd activation results in coordinate upregulation of VE-Cadherin, indicating enhanced endothelial junction formation and vascular stabilization (n = 3 independent experiments). Scale bars, 50 μm. **(C);** Left channel: Western blot analysis of human brain microvascular endothelial cells (HBMECs) infected with pLOC lentivirus expressing human PRND or pLOC empty vector control (GFP/RFP). Prnd overexpression increases intracellular VEGF levels and decreases angiostatin (an endogenous angiogenesis inhibitor), demonstrating that Prnd shifts the angiogenic balance toward pro-angiogenic signaling (n = 3 independent experiments). Right channel: Western blot analysis showing phosphorylation status of key signaling molecules in HBMECs following Prnd overexpression. Enhanced phosphorylation of focal adhesion kinase (FAK), protein kinase B (Akt), extracellular signal-regulated kinases 1/2 (Erk1/2), and SRC family kinases demonstrates that Prnd activates multiple pro-angiogenic signaling pathways, including those regulating cell survival, migration, proliferation, and vascular remodeling (n = 3 independent experiments).

## References

1. Tiedt, S., et al., The neurovascular unit and systemic biology in stroke - implications for translation and treatment. Nature Reviews Neurology, 2022. 18(10): p. 597–612.

2. Makin, S.D., et al., Cognitive impairment after lacunar stroke: systematic review and meta-analysis of incidence, prevalence and comparison with other stroke subtypes. J Neurol Neurosurg Psychiatry, 2013. 84(8): p. 893–900.

3. Savva, G.M., B.C. Stephan, and G. Alzheimer’s Society Vascular Dementia Systematic Review, *Epidemiological studies of the effect of stroke on incident dementia: a systematic review*. Stroke, 2010. 41(1): p. e41–6.

4. Dong, T., et al., Construction and imaging of a neurovascular unit model. Neural Regen Res, 2022. 17(8): p. 1685–1694.

5. McConnell, H.L., et al., The Translational Significance of the Neurovascular Unit. J Biol Chem, 2017. 292(3): p. 762–770.

6. Stamatovic, S.M., et al., Involvement of Epigenetic Mechanisms and Non-coding RNAs in Blood-Brain Barrier and Neurovascular Unit Injury and Recovery After Stroke. Front Neurosci, 2019. 13: p. 864.

7. Sladojevic, N., et al., Claudin-1-Dependent Destabilization of the Blood-Brain Barrier in Chronic Stroke. J Neurosci, 2019. 39(4): p. 743–757.

8. Yang, C., et al., Neuroinflammatory mechanisms of blood-brain barrier damage in ischemic stroke. Am J Physiol Cell Physiol, 2019. 316(2): p. C135–C153.

9. Yang, Y. and M.T. Torbey, Angiogenesis and Blood-Brain Barrier Permeability in Vascular Remodeling after Stroke. Curr Neuropharmacol, 2020. 18(12): p. 1250–1265.

10. Ciric, D. and H. Rezaei, Biochemical insight into the prion protein family. Front Cell Dev Biol, 2015. 3: p. 5.

11. Didonna, A., Prion protein and its role in signal transduction. Cell Mol Biol Lett, 2013. 18(2): p. 209–30.

12. Chen, Z., et al., The vascular endothelial cell-expressed prion protein doppel promotes angiogenesis and blood-brain barrier development. Development, 2020. 147(18).

13. Shyu, W.C., et al., Overexpression of PrPC by adenovirus-mediated gene targeting reduces ischemic injury in a stroke rat model. J Neurosci, 2005. 25(39): p. 8967–77.

14. Li, A., et al., Physiological expression of the gene for PrP-like protein, PrPLP/Dpl, by brain endothelial cells and its ectopic expression in neurons of PrP-deficient mice ataxic due to Purkinje cell degeneration. Am J Pathol, 2000. 157(5): p. 1447–52.

15. Tamguney, G., et al., Genes contributing to prion pathogenesis. J Gen Virol, 2008. 89(Pt 7): p. 1777–1788.

16. Fang, W., et al., CCR2-dependent monocytes/macrophages exacerbate acute brain injury but promote functional recovery after ischemic stroke in mice. Theranostics, 2018. 8(13): p. 3530–3543.

17. Wang, T., et al., GPR68 Is a Neuroprotective Proton Receptor in Brain Ischemia. Stroke, 2020. 51(12): p. 3690–3700.

18. Wang, T., M. He, and X.M. Zha, Time-dependent progression of hemorrhagic transformation after transient ischemia and its association with GPR68-dependent protection. Brain Hemorrhages, 2020. 1(4): p. 185–191.

19. Longa, E.Z., et al., Reversible middle cerebral artery occlusion without craniectomy in rats. Stroke, 1989. 20(1): p. 84–91.

20. Zudaire, E., et al., A computational tool for quantitative analysis of vascular networks. PLoS One, 2011. 6(11): p. e27385.

21. Morita, K., et al., Endothelial claudin: claudin-5/TMVCF constitutes tight junction strands in endothelial cells. J Cell Biol, 1999. 147(1): p. 185–94.

22. Van Itallie, C.M., et al., ZO-1 stabilizes the tight junction solute barrier through coupling to the perijunctional cytoskeleton. Mol Biol Cell, 2009. 20(17): p. 3930–40.

23. Haley, M.J. and C.B. Lawrence, The blood-brain barrier after stroke: Structural studies and the role of transcytotic vesicles. J Cereb Blood Flow Metab, 2017. 37(2): p. 456–470.

24. De, A., et al., The β8 integrin cytoplasmic domain activates extracellular matrix adhesion to promote brain neurovascular development. Development, 2022. 149(6).

25. Claxton, S., et al., Efficient, inducible Cre-recombinase activation in vascular endothelium. Genesis, 2008. 46(2): p. 74–80.

26. Halder, S.K., A. Sapkota, and R. Milner, The importance of laminin at the blood-brain barrier. Neural Regen Res, 2023. 18(12): p. 2557–2563.

27. Sugiyama, S., et al., The tight junction protein occludin modulates blood-brain barrier integrity and neurological function after ischemic stroke in mice. Sci Rep, 2023. 13(1): p. 2892.

28. Selim, M.S., et al., Claudin 5 Across the Vascular Landscape: From Blood-Tissue Barrier Regulation to Disease Mechanisms. Cells, 2025. 14(17).

29. Hatakeyama, M., I. Ninomiya, and M. Kanazawa, Angiogenesis and neuronal remodeling after ischemic stroke. Neural Regen Res, 2020. 15(1): p. 16–19.

30. Zhu, H., et al., Inflammation-Mediated Angiogenesis in Ischemic Stroke. Front Cell Neurosci, 2021. 15: p. 652647.

31. Hu, B., et al., Mechanisms of Postischemic Stroke Angiogenesis: A Multifaceted Approach. J Inflamm Res, 2024. 17: p. 4625–4646.

32. Setyopranoto, I., et al., Comparison of Mean VEGF-A Expression Between Acute Ischemic Stroke Patients and Non-Ischemic Stroke Subjects. Open Access Maced J Med Sci, 2019. 7(5): p. 747–751.

33. Bernardo-Castro, S., et al., Pathophysiology of Blood-Brain Barrier Permeability Throughout the Different Stages of Ischemic Stroke and Its Implication on Hemorrhagic Transformation and Recovery. Front Neurol, 2020. 11: p. 594672.

34. Nguyen, J.N., et al., CD13 facilitates immune cell migration and aggravates acute injury but promotes chronic post-stroke recovery. J Neuroinflammation, 2023. 20(1): p. 232.

35. Shah, F.H., et al., Targeting vascular endothelial growth receptor-2 (VEGFR-2): structural biology, functional insights, and therapeutic resistance. Arch Pharm Res, 2025. 48(5): p. 404–425.

36. Shibuya, M., Vascular Endothelial Growth Factor (VEGF) and Its Receptor (VEGFR) Signaling in Angiogenesis: A Crucial Target for Anti- and Pro-Angiogenic Therapies. Genes Cancer, 2011. 2(12): p. 1097–105.

37. Wilkinson, R., et al., Platelet endothelial cell adhesion molecule-1 (PECAM-1/CD31) acts as a regulator of B-cell development, B-cell antigen receptor (BCR)-mediated activation, and autoimmune disease. Blood, 2002. 100(1): p. 184–93.

38. Newman, P.J., Switched at birth: a new family for PECAM-1. J Clin Invest, 1999. 103(1): p. 5–9.

